# Dissecting Tissue Architecture and Function through Mapping of Cellular Communication Networks

**DOI:** 10.1101/2025.06.21.660290

**Authors:** Dongyu Li, Ming Yang, Lin Hou

## Abstract

Spatial transcriptome provides critical insights for inferring cell-cell communications. Based on the heuristic that physically adjacent cells are more likely to communicate with each other, we develop a computational framework that Infers the Cell-Cell Communication network (IC3) and subsequently identifies communication hotspots with rigorous error control. We demonstrate in simulations and real datasets that our method outperforms existing methods in accuracy and especially improves sensitivity to identify communications involving rare cell types. Applying IC3 in a mouse brain dataset, we recovered how cells communicate with each other to form a local structure that regulates the balance of the blood-brain barrier. These findings highlight that our method is effective in revealing tissue architecture and function from cellular communication networks.

---

Cell-cell communication is essential for the function of multi-cellular organisms [1]. Cells frequently communicate via the specific binding of a ligand secreted by one cell to a receptor of another cell. Such ligand receptor (LR) interactions set in motion a cascade of downstream signaling events, which regulate the activity of transcription factors and lead to subsequent changes in gene expression profiles [2]. Recent advances in single-cell RNA-seq experiments have facilitated the study of cell-cell communication by scoring the expression levels of LR pairs [3, 4]. Supplementing gene expression profiles, spatial transcriptomics data (ST) [5–9] reveal the spatial coordinates of single cells, thus providing an adjacency matrix of cells. Several computational approaches [4, 10–18] have been developed to infer cell-cell communication from spatial transcriptomic data, with the heuristics that neighboring cells with coordinated expression pattern of LR pairs are more likely to communicate with each other.

Existing methods are effective in identifying cell-cell communications at the cell-type level. However, heterogeneity among cells of the same cell type is overlooked. It is not unreasonable to think that cells of the same cell type may behave differently with respect to their communication status. Investigating the pattern of heterogeneous communication can provide a deeper biological understanding. Toward the goal of deciphering the heterogeneous cellular network, we present IC3, a probabilistic graph model that Infers Cell-Cell Communication with single-cell resolution from ST data. Our model identifies the most likely configuration of the cellular communication network, and distinguishes the communication status for cells of the same type. Given the cellular communication network, we further investigate whether or not the differential communication behavior has spatial patterns. We have also developed a scan algorithm to identify hotspots in cell-cell communications and test their significance.

We build a pairwise latent state graph model (**Figure S1**) to infer the cell-cell communication network from ST data sets. Our model has two layers. At the observational layer, given a configuration of the latent edges, a pairwise Poisson graphical model [19] is used to characterize the co-expression pattern of LR in communicating cells. For the latent layer, the probability of network configuration depends on the cell types and the physical distance of the cells. Apparently, our interest lies in inferring the latent status of edges. As the output, IC3 returns the posterior probability of communication between cells. The cellular communication network can facilitate various downstream analyses that improve our understanding of the coordinating mechanism of multicellular organisms.

In the following, we show with real data results and simulations that IC3 identifies interactions at the cell type level with higher accuracy compared to existing methods. The precision of the inference results at the single-cell level is verified by the activation of genes that are downstream of the signaling cascades. We also provide two scenarios in which the cellular network provides additional biological insight. First, it enables detection of communication hotspots, which effectively reveal tissue architecture in the mouse brain. Second, association analysis of the cellular network reveals a significant association of communication and cellular neighborhood.

## IC3 accurately detects cell-cell communications in mouse brain

We applied IC3 to a seqFISH+ dataset that studies the subventricular zone (SVZ) and the olfactory bulb (OB) of the mouse brain [8], and compared the performance of IC3 with existing methods [11–16]. The performance of IC3 is stable regarding to varying choice of hyperparameter (**Figure S2**), and ICM-EM algorithm rapidly converged (Supplementary Note 1, **Figure S3**). In order to create a gold standard dataset of experimentally validated cell-cell communications, we sourced data from both the citeDB database [20], and also manually curated a list from the PubMed literature, including 29 and 24 communications for the SVZ and OB regions (see Table S1 and S2). Note that the detection accuracy was evaluated at the cell-type level. For existing methods, a communication score was calculated for every pair of cell types (see Methods for details). For IC3, we calculated the posterior probability of communication between two types of cells and used network diagrams for visualization (**Figure 1a-b**). IC3 achieved the best AUC in both the SVZ and OB regions (**Figure 1c-d**). IC3 accurately recaptured the cell-cell communications previously reported in the literature and revealed the molecular mechanism underlying the communication. For example, in SVZ, IC3 identified a high posterior probability of communication between neural stem cells and neural progenitor cells, which is documented in existing literature [21]. In OB, IC3 revealed a high posterior probability of communication between astrocytes and olfactory ensheathing cells (**Figure 1b**). Specifically, olfactory ensheathing cells are associated with distinct stress responses in astrocytes, such as boundary formation and increased glial fibrillary acidic protein expression, underscoring their unique role in modulating astrocyte interactions [22].

**Fig. 1:**
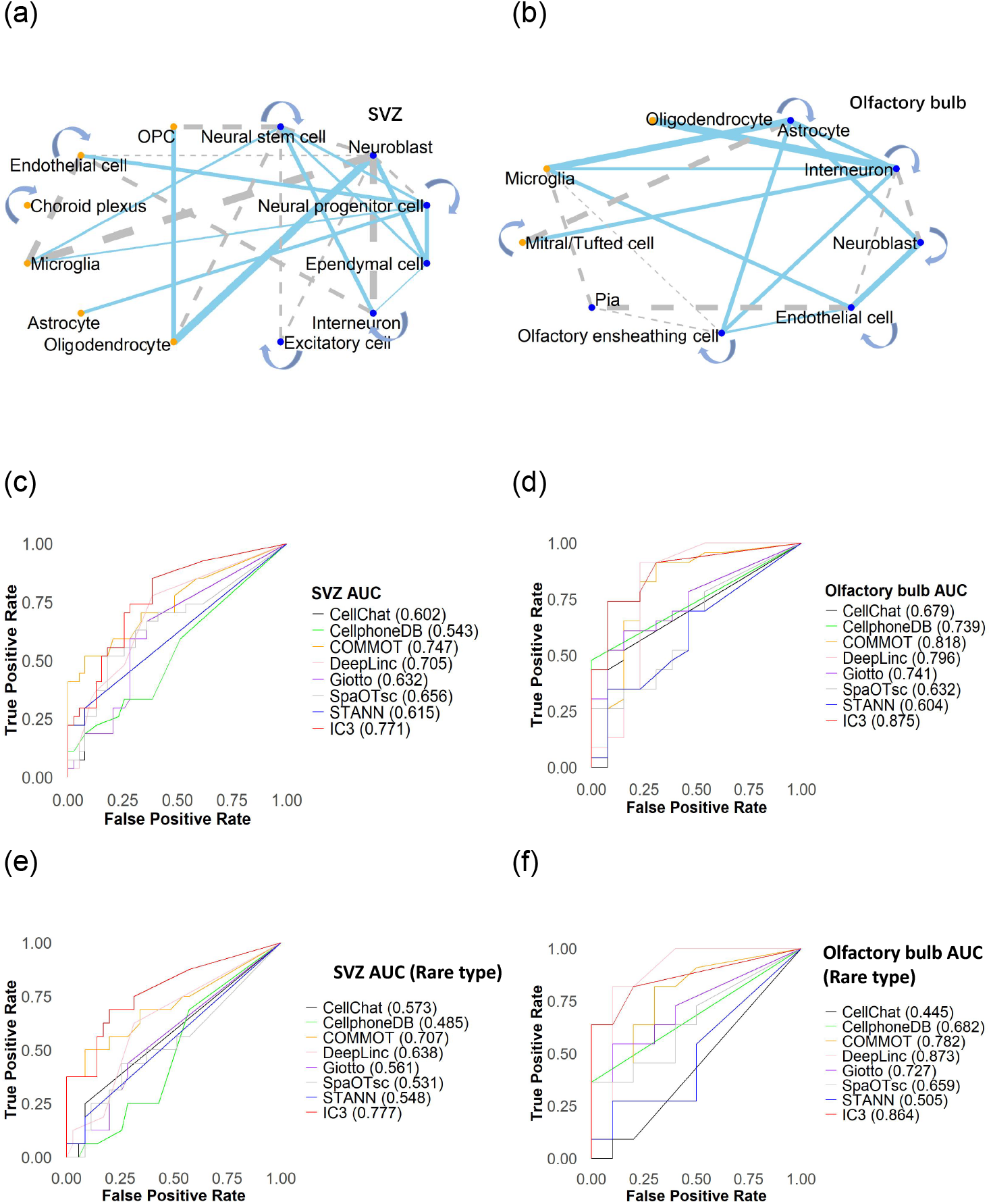
Application of IC3 to the seqFISH+ dataset of the subventricular zone (SVZ) and the olfactory bulb region in mouse brain. (a,b): Communication probabilities between cell type pairs, each dot represents a cell type. Orange dots represent rare cell types, the thickness of the connecting line represents the communication probability, the blue solid line represents that the communication between cell types is supported by the literature, and the gray dashed line indicates that there is currently no literature support. OPC: Oligodendrocyte Precursor Cell. SVZ: (a) Olfactory bulb: (b). (c,d) ROC curves and AUC values for different methods predicting cell-type level communication. SVZ: (c) Olfactory bulb: (d). (e, f) ROC curves and AUC values for different methods predicting cell-type level communication involving at least one rare cell type. SVZ: (e) Olfactory bulb: (f).

Next, we benchmarked IC3 against existing methods for capturing communications that involve at least one rare cell type. Despite individual rarity, these populations collectively constitute a significant portion of cells. For example, in the mouse brain SVZ dataset, the number of cells of rare cell types (no more than 10 cells) sums up to 43 out of 281, and the estimated number of communications is 21 out of 97. IC3 generally outperformed the other methods in AUC values (**Figure 1e-f**). In OB, the AUC of DeepLinc is slightly higher than IC3. In addition to the advantage in AUC values, IC3 identified cell-cell communications that are ignored by the other methods. For example, IC3 identified a high probability of communication between neuroblasts and oligodendrocytes in SVZ. The existing literature supports the involvement of neuroblasts in the generation and communication of oligodendrocytes in the adult mouse brain [23, 24]. In OB, IC3 detected communication between mitral/tufted cells with interneuron. Previous studies have shown that interneurons can establish dendritic contacts with mitral/tufted cells during their growth process [25].

To further demonstrate that IC3 is more accurate in inferring communications of rare cell types, we simulated ST data at various levels of cell type frequency (see Supplementary Note 2) to evaluate IC3 and existing methods. We found that all methods have high AUC values (*AUC >* 0.85) for frequently observed cell types (**Figure S6a**). For rare cell types, the accuracy decreased when the frequency of the cell type under consideration decreased, and IC3 outperformed the other methods when the frequency of the rare cell type was between 5% to 10% (**Figure S6b**).

## Communication hotspots reveals tissue microstructure in the mouse brain

The cellular communication map revealed by IC3 provides a unique opportunity to investigate the spatial clustering pattern of cell-cell communications and hence its functional relevance. For this purpose, we develop a scan algorithm (see Methods for details) to identify communication hotspots, which are local areas with concentrated cell-cell communications. We applied IC3 to a MERFISH dataset of the hypothalamic preoptic region of mouse brain [26]. The dataset is consisted of 10 mouse samples, and each mouse sample has multiple slices. For brevity, we only illustrate the results of mouse 1, which has the most number of slices, since the extensive tissue section is particularly well-suited for analyzing spatial heterogeneity. We scanned the communication network to identify hotspots (**Figure S7-18**). The significant results readily recovered known tissue architecture.

There were several communication hotspots foretween pericytes and endothelial cells (Peri-Endo hotspot). After aligning the coordinates of different slices, we found that the Peri-Endo hotspots in several slices form a perpendicular three-dimensional structure (**Figure 2a**). The role of cellular communication between endothelial cells and pericytes is essential for the formation, stability, and function of blood vessels [27]. In particular, pericytes and endothelial cells physically connect to each other to form a functional unit, which is crucial to regulate the blood-brain barrier in the hypothalamic region [28, 29]. The diameter of the Peri-Endo hotspot is around 100 microns, which is about the typical size of blood vessels in the mouse brain. The results highlight the connection between communication hotspot and tissue microstructure.

**Fig. 2:**
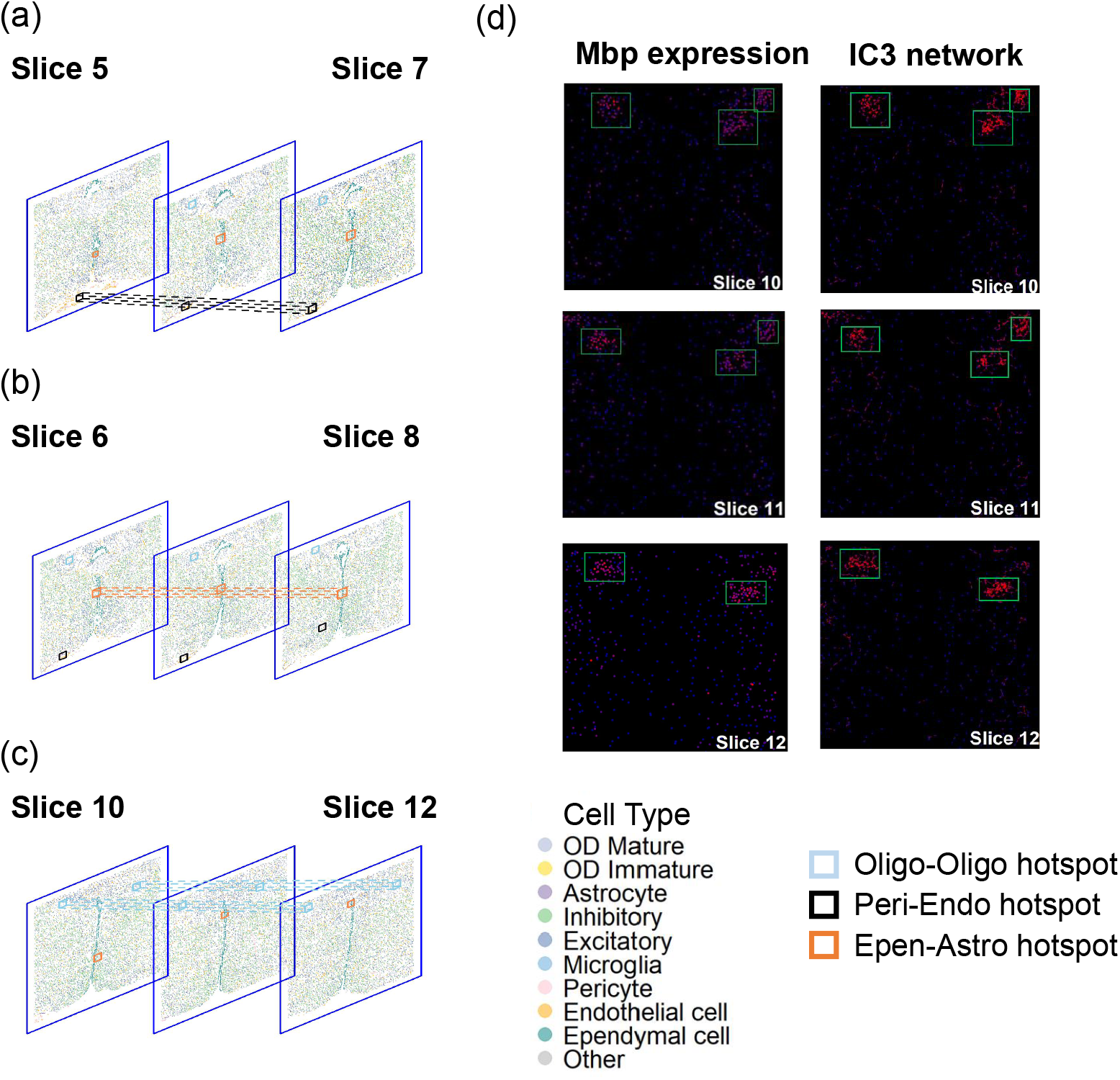
IC3 reveals spatial clustering effect of cell-cell communications in a MERFISH dataset of mouse brain. The cell-level communication network for each slice is calculated to identify communication hotspots using scan statistics. If hotspots in adjacent slices are closely positioned, they are combined to form a three-dimensional communication hotspot region. (a) Communication hotspot region in slice 5-7 of mouse 1. (b) Communication hotspot region in slice 6-8 of mouse 1. (c) Communication hotspot region in slice 10-12 of mouse 1. (d) The first column shows a heatmap of Mbp gene (myelin-related gene) expression for three slices, with green boxes highlighting hotspot regions. The second columns display network diagrams that infer the communication strength between oligodendrocytes using IC3. Oligo-Oligo hotspot: Communication hotspot between oligodendrocytes. Peri-Endo hotspot: Communication hotspot between pericytes and endothelial cells. Epen-Astro hotspot: Communication hotspot between ependymal cells and astrocytes.

Similarly, communication hotspots of ependymal cells and astrocytes (Epen-Astro hotspot) in adjacent slices form three-dimensional communication hotspots around the periventricular hypothalamic nucleus in the preoptic region (**Figure 2b**). Ependymal cells play a vital role in regulating the function of the central nervous system, including the production and circulation of cerebrospinal fluid, brain metabolism and waste removal, and this process is related to the communication with astrocytes [30, 31]. Astrocytes regulate central nervous function by maintaining the integrity of the cerebrospinal fluid-brain barrier [32]. Thus, the Epen-Astro hotspot may play a critical role in facilitating cerebrospinal fluid dynamics within the preoptic area of the hypothalamus and in regulating central nervous system function.

Additionally, we observed that oligodendrocyte communication was concentrated within three-dimensional regions which aligned well with the structure of the fornix–major myelinated fiber tracts of the rostral hypothalamus (**Figure 2c**). Oligodendrocytes and the communications between them are essential for the process of myelination. In the nervous system, myelin proteins exhibit varying degrees of aggregation, contributing to distinct functional and structural roles [33, 34]. MBP and PLP are the major proteins in central nervous system myelin and play essential roles in compaction and stabilization of the myelin sheath [35]. Consistently, we found that the expression of Mbp gene was abundant in the Oligo-Oligo hotspot (**Figure 2d, Figure S19**). The combined evidence implicates that the identified hotspot are closely related to the process of myelination of the fornix–major myelinated fiber tracts of the rostral hypothalamus.

In summary, from the fine-resolution communication network provided by IC3, one can distinguish fragmentary communications and spatially clustered communications. We showed that clustered communications, or communication hotspot effectively recover known tissue microstructure. Subsequently, we expect that identification of communication hotspot can generate biologically plausible hypotheses of tissue microstructure and function.

## IC3 distinguishes communication status in a small neighborhood

Oftentimes, when a cell of type A communicates with a cell of type B, there are more than one cells of type B in its neighborhood. Below we show that with IC3 output, we can precisely tell which cell is engaged in the communication. We use a toy simulation model to illustrate the advantage of system modeling in IC3. Consider a system of three cells, one of type A sending out ligand *L*, and two of type B expressing *R*_1_ and *R*_2_ as the receptors respectively. In the simulation, we fixed the expression level of *L*, varied the expression levels of *R*_1_ and *R*_2_, and ran IC3 to estimate the latent communication status *W*_1_ and *W*_2_ (see Supplementary Note 2). The results changed with varying levels of the receptors, as expected (**Figure 3a**). In detail, when *R*_1_ is expressed at insufficient levels and *R*_2_ is abundant, the cell of type A communicates exclusively with the cell expressing *R*_2_. When both *R*_1_ and *R*_2_ are abundant, *W*_1_ = 1, *W*_2_ = 1 gives the maximum posterior probability. The toy example illustrates the advantage of systematic modeling of the IC3 model, that it adaptively learns from the local neighborhood to precisely determine the communication status of individual cells. In other words, the model makes inferences with awareness of all adjacent cells. In contrast, pseudo-bulk methods, which sum over cells of the same cell type and make inference at cell type level, cannot enjoy such precision by principle.

**Fig. 3:**
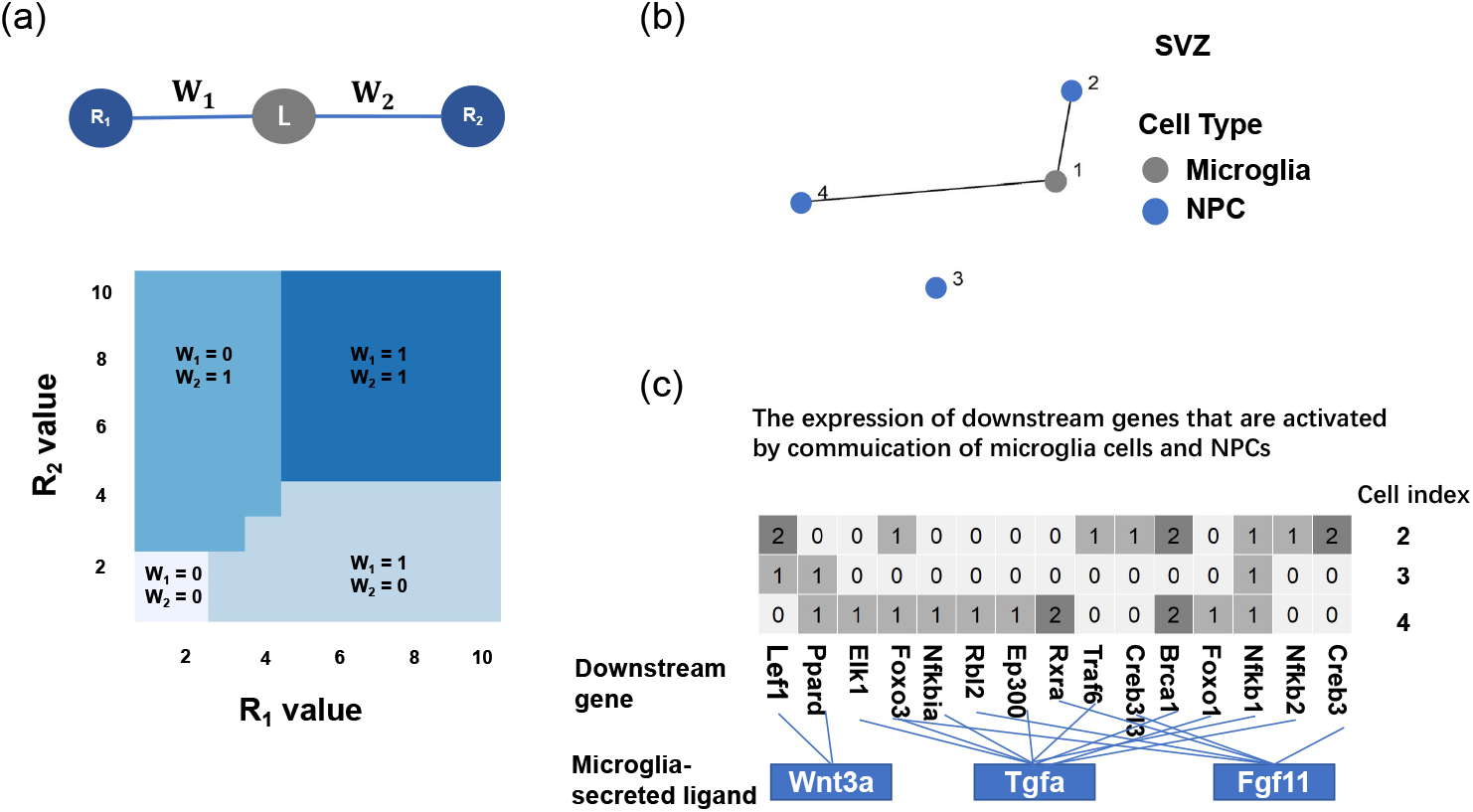
(a) The toy model consists of three cells. Cell expressing the ligand has a cell expressing the receptor on each side. The color map below represents the relationship between the IC3 model’s posterior estimate of the cell level communication status W and the expression value of the cell receptors on both sides (*R*_1_, *R*_2_). (b): A small area in SVZ containing several cells. Colors represent cell types, and lines indicate posterior probability of communication between pairs of cells greater than 0.5. (c) Heatmap depicting expression profiles of downstream target genes in neural progenitor cells (NPCs) regulated by microglia-secreted ligands. A green annotation bar at the bottom denotes microglia-derived ligand genes, while connecting lines indicate regulatory relationships between these ligands and their downstream targets in NPCs.

We further validate IC3 by examining the expression level of genes that are downstream to the communication signaling event in real datasets. From the SVZ slice in the seqFISH+ dataset, we selected a specific region defined by x-coordinates between 890 and 120 and y-coordinates between 940 and 1140, containing 3 neural progenitor cells (NPCs) and 1 microglia (**Figure 3b**). IC3 identified communication between the microglia cell and the NPC cells 2 and 4. Communication between microglia and neural progenitor cells (NPCs) has previously been reported in the central nervous system [36, 37].

Subsequently, we checked the expression levels of downstream genes in NPCs that are regulated by microglia-secreted ligands (**Figure 3c**). The expression levels of downstream genes were higher in Cell 2 and Cell 4 compared to Cell 3, which echoed the predictions of IC3 regarding the different communication status of cells. Note that the expression levels of downstream genes are not considered in the IC3 model, thus their differential expression pattern provides independent verification of the IC3 results. When we compared the dropout rate and average gene expression in communicating and non-communicating cells, no significant difference was observed (**Figure S20**).

## Local neighborhood significantly affects cell-cell communications

With the observation that pairs of cells of the same type composition and similar distance exhibit differential communication status, we explored potential influential factors. In particular, we hypothesize that the local neighborhood, the presence or absence of a third cell type in close proximity, may contribute. We designed a contingency table test for the association. Specifically, for a triplet of cell types (A, B, C) (A, B, and C could be identical), we tested whether the presence of a cell of type C in the neighborhood of cells of type A and B would affect the communication status between A and B. A contingency table was constructed (see Methods, **Figure 4a**). We enumerated all possible cell type triplets across the 12 slices of mouse 1 in the MERFISH dataset, and identified a few sigficant triplets (q values*<*0.05). For example, the presence of astrocytes enhances the communication between inhibitory and excitatory (**Figure 4b**), which is consistent with previous reports that astrocytes facilitate the formation and function of excitatory and inhibitory synapses in the central nervous system [38]. We also find that the communication between oligodendrocytes were significantly correlated with the presence of ependymal cells in the anterior commissure region (**Figure 4b**). The findings highlight that the composition of local neighborhood plays an important role in cell-cell communications.

**Fig. 4:**
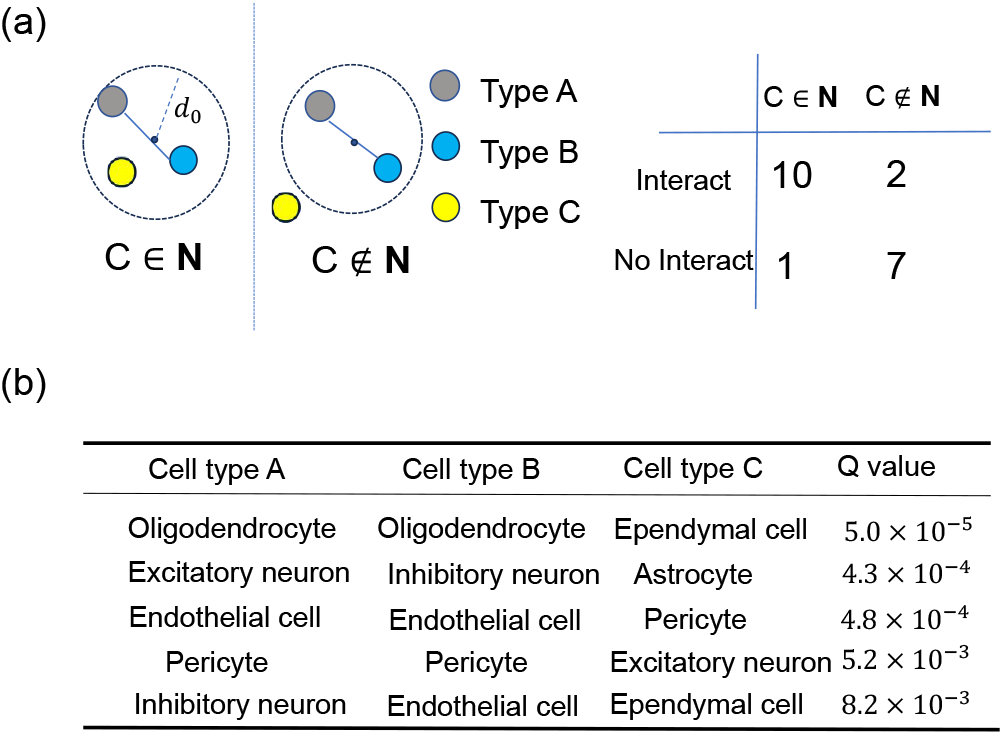
Contingency table analysis of cellular communication network. (a) Test for association between the communication status between cell types A and B and their neighbors. N denotes the set of cells within the *d*_0_ neighborhood of the pair of cells in consideration. (b) The list of cell type triplets with q value*<*0.05.

## Discussion

We present IC3, a probabilistic approach for inferring cell-cell communication from spatial transcriptomic data. The output of IC3 characterizes cellular communications at multiple levels. It estimates the probability of communication between two cell types, and infers the latent communication status between two cells. We further deploy a scan algorithm to identify communication hotspots from the IC3 outputs. Evaluated on various experimental platforms, IC3 is more accurate in identifying cell-cell communications than existing methods. More importantly, IC3 has two unique advantages. First, it provides a single cell resolution communication network, facilitating the study of the heterogeneity of cellular communications. For example, the scan approach we developed recovers key cellular communications to maintain the blood-brain barrier, showcasing its ability to precisely reveal the functional architecture of tissues. Second, IC3 jointly models all cells and their connections in a study, hence provides a panoramic view of the communication network. In contrast, existing methods treat each cell and each communication separately, and to sum up the fragmented predictions of cell-cell communications not necessarily leads to a reasonable estimate of the entire network (**Figure 3**).

While IC3 offers significant advantages over existing methods, it has several limitations. First, our current implementation only support spatial transcriptomic data with single cell resolution. An extension is under development to accommodate spot-level data. Second, the current model relies solely on the expression level of ligands and receptors, while we anticipate that incorporating other genes in relevant signaling pathways in the model could potentially improve the performance of the method.

Looking ahead, IC3 holds promise for several key applications. It can be extended to study differential communication across disease states, revealing how disruptions in cell-cell communication may contribute to pathogenesis. Additionally, by focusing on communication hotspots, we could align adjacent spatial transcriptomic slices, creating more comprehensive tissue maps that better reflect the functional and structural organization of tissues.

## Methods

### Gold standard dataset of cell-cell communications

For each tissue, we created a gold standard list of cell-cell communications by querying the PubMed literature and CiteDB database [20]. Specifically, for each pair of cell types, we searched in PubMed using the names of the two cell types combined with the keywords “interact” or “communicate” from the retrieved literature, we manually evaluated whether the studied tissue matched the tissue of interest and whether the retrieved text explicitly documented experimental evidence of cell-cell communication between the two cell types. Only pairs that met the above criteria were included in the gold standard dataset. We also included all communications annotated in the context of brain. For more specific brain tissues. The curated data is provided in **Table S1-S3**.

### Logistic model for latent communication status

IC3 builds a probabilistic model of cell-cell communication, while the communication status is a binary latent variable. Consider an ST data set with *N* cells of *T* cell types and *G* genes. Note that only genes that encode ligands and receptors are considered. Let **Y** be the gene expression matrix, of dimensions *N × G*. **Y**_*cg*_ is the read count of the gene *g* of the cell *c*. **D** is the distance matrix between cells, calculated from their location coordinates. At the cell type level, **K** is the matrix of probability of communication, as **K**_*st*_ denotes the probability of communication between cell types *s* and *t*. We further transform **K** into 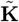, such that 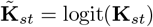. At the individual cell level, **W** is the matrix of latent communication status, where 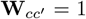 if the cells *c* and 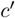 are communicating and 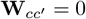 otherwise.

For each pair of cells, the probability of communication is determined by the cell types involved and the physical distance between them. In particular, for two cells *c* and *c*^′^ with distance less than *d*_0_, the probability of communication is determined by 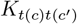 and their distance 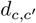, where *t*(*c*) and *t*(*c*^′^) are the respective cell types, by logistic regression (Equation 1). For cells with distance greater than *d*_0_, we set the probability of communication to zero.

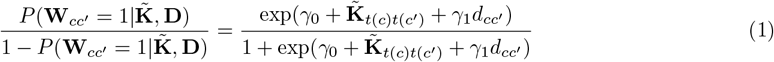

*γ*_0_ and *γ*_1_ are the linear coefficients. We did not include a coefficient for 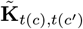 to keep the parameters identifiable.

### Pairwise Poisson graphical model

An undirected graph (*V, E*) is constructed from ST data and the latent communication network, where each vertex (*c, g*) denotes the expression of gene *g* of cell *c*. Two vertices are connected if the corresponding cells are communicating and the corresponding genes form LR pairs. We use the Pairwise Poisson Graphical Model [19] to characterize the co-expression pattern of LR pairs in communicating cells (see Equation 2).

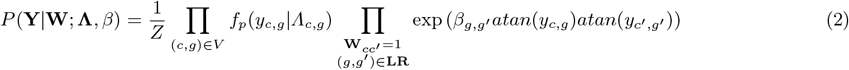

*f*_*p*_(*·*|*λ*) denotes the density function of Poisson distribution with mean value *λ*. *Z* is the partition function. We transform the raw read counts *y*_*c,g*_ by the *atan* function so that *Z* will not go to infinity. *β* is a vector of graphical parameters, reflecting the communication strength of the ligand *g* and the receptor *g*^′^. For each node, the Poisson parameters is the product of the total reads count of cell *c* and a cell type specific parameter of the gene, *Λ*_*c,g*_ = *n*_*c*_ *× λ*_*t*(*c*),*g*_ [39].

We estimate all parameters *Θ* = (**Λ**, *γ*_0_, *γ*_1_, *β*, **K**) and the latent state **W** by maximizing the following log likelihood function:

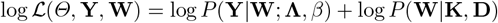

We further approximated the likelihood function by the pseudo-likelihood function (Equation 3), and used the ICM-EM algorithm (Supplementary Note 1) for estimation.

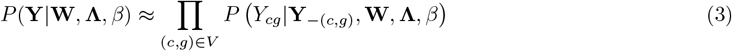

### Scan statistic for detecting communication hotspot

For each slice, we employ scan statistics [40] to identify hotspots in the cellular communication network. Concerning communications between cells of type A and B, we take square windows of varying sizes and calculate the likelihood ratio test (LRT) statistic (Equation 4) that evaluates the enrichment of cellular communications in the window. In particular, the within-window proportion of communicating neighbors is calculated as

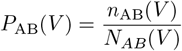

where *N* ^AB^(*V*) is the number of cell pairs of type A and B with distance less than *d*_0_ in the window *V*, and *n*^AB^(*V*) is the number of cell pairs of type A and B with posterior communication probability greater than 0.5. Similarly, the background proportion of communicating neighbors is calculated as

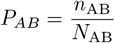

where *N*_AB_ is the total number of cell pairs of type A and B with distance less than *d*_0_ in the slice, and *n*_AB_ is the corresponding number of cell pairs of type A and B with posterior communication probability greater than 0.5.

The corresponding LRT statistic is

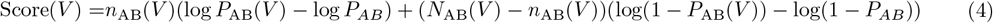

We enumerate windows of varying sizes and locations to identify the optimal score and evaluate its significance by permutation test (Supplementary Note 3).

### Contingency table analysis of cellular communication

All pairs of cells satisfying the following condition are examined: (1) one of the cells is of type A and the other is of type B and (2) the distance between them is less than *d*_0_. Then we cross-tabulate the cell pairs by two factors (Fig 4a). Considering a pair of cells *c* and *c*^′^, the first factor is whether or not *c* and *c*^′^ have communication 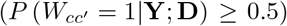. The second factor is the existence of any cells of type C within *d*_0_ distance from the midpoint of the location of *c* and *c*^′^. We use Fisher’s exact test to test whether the existence of cells of type C affects the communication between cells of type A and B, and control the false discover rate by the Benjamini-Hochberg method.

### Comparison between different Methods

The output differs among the compared methods, and to facilitate the comparison, we define communication strength of a pair of cell types A and B for each method. For IC3, the estimated communication probability between two cell types is used. For CellChat, CellphoneDB, Giotto, and STANN, we take the number of significant LR pairs as the communicating strength between cell type *A* and *B* [20]:

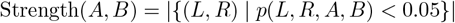

where *p*(*L, R, A, B*) denotes the adjusted p-value for the association between LR pair (*L, R*) and cell type pair (*A, B*). For SpaOTsc, COMMOT, and DeepLinc, the communication strength between cell types *A* and *B* is the average over all cell-level interaction strengths:

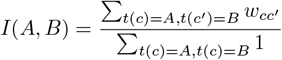

where 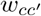 is the communication strength between cells *c* and *c*^′^ as provided by the corresponding software.

## Supporting information

Supplementary for IC3

## Data availability

The seqFISH+ dataset was obtained from https://github.com/CaiGroup/seqFISH-PLUS. The MERFISH dataset of the hypothalamic preoptic region was obtained from https://datadryad.org/dataset/doi:10.5061/dryad.8t8s248.

## Code availability

The open-source software is available at https://github.com/lidongyu16/IC3.

## Acknowledgements

We acknowledge research support from the National Natural Science Foundation of China (Grant No. T2322017).

