## Supplementary for IC3 for "Dissecting Tissue Architecture and Function through Mapping of Cellular Communication Networks"

June 18, 2025

**3 Supplementary Note 3: The scan statistic and permutation test** **8**

**4 Supplementary Tables** **10**

**5 Supplementary Figures** **12**

IC3 is a probabilistic model designed to infer cell-cell communication from spatial transcriptomic data, where the communication status between cell pairs is a binary latent variable. The model integrates information on gene expression, spatial distance, and cell type information.

Suppose $N$ is the number of cells, $T$ is the number of cell types, and $G$ is the number of ligands/receptors. The notations are listed below.

- $\mathbf{Y} \in \mathbb{R}^{N\times G}$: the gene expression matrix.
- $\mathbf{D} \in \mathbb{R}^{N\times N}$: the distance matrix between cells.
- $\mathbf{K} \in [0,1]^{T\times T}$: the matrix of probability of communication.
- $\tilde{\mathbf{K}} = \text{logit}(\mathbf{K})$: logit-transformation of $\mathbf{K}$.
- $\mathbf{W} \in \{0,1\}^{N\times N}$: the matrix of latent binary communication status.
- $\mathbf{\Theta} := \{\beta, \lambda, \gamma, \mathbf{K}\}$ The set of all parameters.

In the following, we describe the ICM-EM algorithm [1] used for parameter estimation in detail.

### 1 Supplementary Note 1: Details of the ICM-EM algorithm

The **ICM-EM algorithm** iterates through three key steps:

1. **ICM Step**: Computes the expectation of latent variables $e_{c_1c_2} := \mathbb{E}\left[W_{c_1c_2}|\mathbf{Y};\Theta,\hat{\mathbf{W}},\mathbf{D}\right]$ given previous estimates while recording their MAP (maximum a posteriori) solutions.
2. **E Step**: Constructs the Q function:
$$Q(\Theta|\Theta^{(t)}) = \mathbb{E}_{\mathbf{W}|\mathbf{Y},\Theta^{(t)}}[\log\mathcal{L}(\mathbf{Y},\mathbf{W}|\Theta)]$$
where $\log\mathcal{L}(\mathbf{Y},\mathbf{W}|\Theta)$ is the log-likelihood of complete data. The ICM outputs yield a tractable approximation by fixing $\mathbf{W}$ to the MAP estimates.
3. **M Step**: Updates the parameters via $\Theta^{(t)} = \text{argmax}_\Theta\, Q(\Theta|\Theta^{(t-1)})$

Suppose that we have done (t-1) iterations and got the parameter $\mathbf{\Theta}^{(t-1)} := \{\beta^{(t-1)}, \lambda^{(t-1)}, \gamma^{(t-1)}, \mathbf{K}^{(t-1)}\}$ and the MAP of latent variables $\mathbf{W}^{(t-1)}$, the following sections will show the algorithm details.

#### 1.1 ICM Step

The ICM step computes the posterior expectation of the latent communication variables $\mathbf{W}$ using the following Bayesian framework.

$$\begin{aligned}
e^{(t)}_{c_1c_2} &:= \mathbb{E}\left[W_{c_1c_2}|\mathbf{Y};\Theta^{(t-1)},\hat{W}^{(t-1)},\mathbf{D}\right]\\
&= P\left(W_{c_1c_2}=1|\mathbf{Y};\Theta^{(t-1)},\hat{W}^{(t-1)},\mathbf{D}\right)\\
&= \frac{P\left(W_{c_1c_2}=1,\mathbf{Y}|\Theta^{(t-1)},\hat{W}^{(t-1)},\mathbf{D}\right)}{P\left(W_{c_1c_2}=1,\mathbf{Y}|\Theta^{(t-1)},\hat{W}^{(t-1)},\mathbf{D}\right)+P\left(W_{c_1c_2}=0,\mathbf{Y}|\Theta^{(t-1)},\hat{W}^{(t-1)},\mathbf{D}\right)}\\
&= \frac{b_{c_1c_2}}{1+b_{c_1c_2}}
\end{aligned} \tag{1}$$

where $b_{c_1c_2}$ is defined as:

$$b_{c_1c_2} := \frac{P\left(W_{c_1c_2}=1,\mathbf{Y}|\Theta^{(t-1)},\hat{W}^{(t-1)},\mathbf{D}\right)}{P\left(W_{c_1c_2}=0,\mathbf{Y}|\Theta^{(t-1)},\hat{W}^{(t-1)},\mathbf{D}\right)}$$

By Bayes' formula, we have the following:

$$P(\mathbf{W},\mathbf{Y}|\Theta^{(t-1)},\mathbf{D}) = P(\mathbf{Y}|\mathbf{W};\lambda^{(t-1)},\beta^{(t-1)})\cdot P(\mathbf{W}|\mathbf{K}^{(t-1)};\gamma_0^{(t-1)},\gamma_1^{(t-1)},\mathbf{D})$$

Then $b_{c_1c_2}$ can decompose into two components:

$$b_{c_1c_2} = \underbrace{\frac{P(\mathbf{Y}|W_{c_1c_2}=1,\hat{\mathbf{W}}^{(t-1)},\lambda^{(t-1)},\beta^{(t-1)})}{P(\mathbf{Y}|W_{c_1c_2}=0,\hat{\mathbf{W}}^{(t-1)},\lambda^{(t-1)},\beta^{(t-1)})}}_{b_1} \cdot \underbrace{\frac{P(W_{c_1c_2}=1|\mathbf{K}^{(t-1)},r^{(t-1)},\mathbf{D})}{P(W_{c_1c_2}=0|\mathbf{K}^{(t-1)},r^{(t-1)},\mathbf{D})}}_{b_2} \tag{2}$$

We employ the pseudo-likelihood approximation [2] to simplify $b_1$:

$$P(\mathbf{Y}|W_{c_1c_2}=w,\hat{\mathbf{W}}^{(t-1)},\lambda,\beta) \approx \prod_{(c,g)\in V} P\left(Y_{cg}|\mathbf{Y}_{-(c,g)},W_{c_1c_2}=w,\hat{\mathbf{W}}^{(t-1)},\lambda,\beta\right)$$

The conditional probability for each gene-cell pair is derived as:

$$\begin{aligned} P(Y_{c_0g_0}|\mathbf{Y}_{-(c_0g_0)},\mathbf{W},\lambda,\beta) &= \frac{1}{z_{c_0g_0}(\mathbf{W})} f_{\mathrm{p}}(Y_{c_0g_0}|\Lambda_{c_0g_0}) \\ &\times \prod_{\substack{c'\in\mathcal{N}(c_0)\\ g'\in\mathcal{N}(g_0)}} \exp\left(\beta_{g_0g'}W(c_0,c')\mathrm{atan}(Y_{c_0g_0})\mathrm{atan}(Y_{c'g'})\right) \end{aligned}$$

where the normalizing constant $z_{c_0g_0}(\mathbf{W})$ is:

$$z_{c_0g_0}(\mathbf{W}) = \sum_{y=0}^{\infty} f_{\mathrm{p}}(y|\Lambda_{c_0g_0}) \prod_{\substack{c'\in\mathcal{N}(c_0)\\ g'\in\mathcal{N}(g_0)}} \exp\left(\beta_{g_0g'}W(c_0,c')\mathrm{atan}(y)\mathrm{atan}(Y_{c'g'})\right)$$

Due to the rapid decay of the Poisson distribution, we approximate the infinite series $z_{c_0g_0}(\mathbf{W})$ by truncating at $B = 10$:

$$z_{c_0g_0}(\mathbf{W}) \approx \sum_{y=0}^{B} \frac{\Lambda_{c_0g_0}^{y} e^{-\Lambda_{c_0g_0}}}{y!} \prod_{\substack{c'\in\mathcal{N}(c_0)\\ g'\in\mathcal{N}(g_0)}} \exp\left(\beta_{g_0g'}W(c_0,c')\mathrm{atan}(y)\mathrm{atan}(Y_{c'g'})\right)$$

After the pseudo-likelihood and infinite series approximation with canceling common terms, $b_1$ simplifies to:

$$\begin{aligned} b_1 = \prod_{g} \Bigg[ &\frac{z_{c_1g}(W_{c_1c_2}=1,\hat{\mathbf{W}})z_{c_2g}(W_{c_1c_2}=1,\hat{\mathbf{W}})}{z_{c_1g}(W_{c_1c_2}=0,\hat{\mathbf{W}})z_{c_2g}(W_{c_1c_2}=0,\hat{\mathbf{W}})} \times \\ &\exp\Big(\sum_{g'\in\mathcal{N}(g)} \beta_{gg'}\left(\mathrm{atan}(Y_{c_1g})\mathrm{atan}(Y_{c_2g'}) + \mathrm{atan}(Y_{c_1g'})\mathrm{atan}(Y_{c_2g})\right)\Big)\Bigg] \end{aligned} \tag{3}$$

$b_2$ can be simplified by

$$b_2 = \exp\left(\gamma_0^{(t-1)} + \mathrm{logit}(K_{t(c_1)t(c_2)}^{(t-1)}) + \gamma_1^{(t-1)} d_{c_1c_2}\right) \tag{4}$$

The maximum a posteriori (MAP) estimate is obtained by thresholding:

$$\hat{W}_{c_1c_2}^{(t)} = \begin{cases} 1 & \text{if } e_{c_1c_2}^{(t)} > 0.5 \\ 0 & \text{otherwise} \end{cases} \tag{5}$$

#### 1.2 E-Step

The Q-function representing the expected complete log-likelihood is computed as:

$$\begin{aligned} Q(\Theta|\Theta^{(t-1)}) &= \mathbb{E}_{\mathbf{W}|\mathbf{Y},\Theta^{(t-1)}}\left[\log\mathcal{L}(\mathbf{Y},\mathbf{W}|\Theta)\right] \\ &= \mathbb{E}_{\mathbf{W}}\left[\log P(\mathbf{Y}|\mathbf{W};\lambda^{(t-1)},\beta^{(t-1)}) + \log P(\mathbf{W}|\mathbf{K}^{(t-1)},\gamma^{(t-1)},\mathbf{D})\right] \\ &\approx \underbrace{\mathbb{E}_{\mathbf{W}}\left[\sum_{c,g}\log P(Y_{cg}|\mathbf{Y}_{-(c,g)},\mathbf{W},\lambda,\beta)\right]}_{\text{Pseudo-likelihood expectation}} \\ &\quad + \underbrace{\mathbb{E}_{\mathbf{W}}\left[\sum_{(c_1,c_2)\in\mathbf{CN}}\log P(W_{c_1c_2}|\mathbf{K}^{(t-1)},r^{(t-1)},\mathbf{D})\right]}_{\text{Prior expectation}} \end{aligned}$$

By exchanging the expectation and summation signs, the Q function can be simplified as following:

$$
\begin{aligned}
Q(\Theta|\Theta^{(t-1)}) = &\sum_{c,g}\Bigg[Y_{cg}\log(n_c\lambda_{t(c)g}) - n_c\lambda_{t(c)g} - \log(Y_{cg}!) \\
&+ \sum_{\substack{c'\in\mathcal{N}(c)\\ g'\in\mathcal{N}(g)}} \beta_{gg'}e_{cc'}\arctan(Y_{cg})\arctan(Y_{c'g'}) - \mathbb{E}_{\mathbf{W}}\left[\log z_{cg}(\mathbf{W})\middle|\Theta^{(t-1)}\right]\Bigg] \\
&+ \sum_{(c_1,c_2)\in\mathbf{CN}}\Big[(\gamma_0 + \operatorname{logit}(K_{t(c_1)t(c_2)}) + \gamma_1 d_{c_1c_2})e_{c_1c_2} - \log\big(1+\exp\big(\gamma_0 + \operatorname{logit}(K_{t(c_1)t(c_2)}) + \gamma_1 d_{c_1c_2}\big)\big)\Big]
\end{aligned}
$$

The computationally challenging term $\mathbb{E}_{\mathbf{W}}[\log z_{cg}(\mathbf{W})]$ is approximated using the MAP estimate:

$$
\mathbb{E}_{\mathbf{W}}\left[\log z_{cg}(\mathbf{W})\middle|\Theta^{(t-1)}\right] \approx \log z_{cg}(\hat{\mathbf{W}}^{(t)})
$$

where the partition function $z_{cg}$ is computed via truncated series:

$$
z_{cg}(\hat{\mathbf{W}}^{(t)}) = \sum_{y=0}^{B}\left[\frac{(n_c\lambda_{t(c)g})^y}{y!}e^{-n_c\lambda_{t(c)g}} \times \exp\left(\sum_{\substack{g'\in\mathcal{N}(g)\\ c'\in\mathcal{N}(c)}}\beta_{gg'}\hat{W}_{cc'}^{(t)}\arctan(y)\arctan(Y_{c'g'})\right)\right]
$$

#### 1.3 M-Step

Our final Q function can be written as:

$$
\begin{aligned}
Q(\lambda,\beta,\gamma,K) = &\sum_{c,g}\Bigg[Y_{cg}\log(n_c\lambda_{t(c)g}) - n_c\lambda_{t(c)g} - \log(Y_{cg}!) + \sum_{\substack{c'\in\mathcal{N}(c)\\ g'\in\mathcal{N}(g)}}\beta_{gg'}e_{cc'}\arctan(Y_{cg})\arctan(Y_{c'g'}) \\
&- \log\sum_{y=0}^{B}\Big[\frac{(n_c\lambda_{t(c)g})^y}{y!}e^{-n_c\lambda_{t(c)g}} \times \exp\Big(\sum_{\substack{g'\in\mathcal{N}(g)\\ c'\in\mathcal{N}(c)}}\beta_{gg'}\hat{W}_{cc'}^{(t)}\arctan(y)\arctan(Y_{c'g'})\Big)\Big]\Bigg] \\
&+ \sum_{(c_1,c_2)\in\mathbf{CN}}\Big[(\gamma_0 + \operatorname{logit}(K_{t(c_1)t(c_2)}) + \gamma_1 d_{c_1c_2})e_{c_1c_2} - \log\big(1+\exp\big(\gamma_0 + \operatorname{logit}(K_{t(c_1)t(c_2)}) + \gamma_1 d_{c_1c_2}\big)\big)\Big]
\end{aligned}
$$

Given the absence of closed-form solutions, we employ Newton-Raphson iteration for parameter updates.

**Poisson Rate Parameters $\lambda_{kg}$**

For gene $g$ in cell type $k$:

$$
\begin{aligned}
\frac{\partial Q}{\partial\lambda_{kg}} &= \sum_{c:t(c)=k}\left[\frac{Y_{cg}}{\lambda_{kg}} - n_c\frac{z_{cg}(\hat{\mathbf{W}}^{(t)},1)}{z_{cg}(\hat{\mathbf{W}}^{(t)})}\right] \\
\frac{\partial^2 Q}{\partial\lambda_{kg}^2} &= -\sum_{c:t(c)=k}\left[\frac{Y_{cg}}{\lambda_{kg}^2} + n_c^2\frac{z_{cg}(\hat{\mathbf{W}}^{(t)},2)z_{cg}(\hat{\mathbf{W}}^{(t)}) - z_{cg}(\hat{\mathbf{W}}^{(t)},1)^2}{z_{cg}(\hat{\mathbf{W}}^{(t)})^2}\right]
\end{aligned} \tag{6}
$$

with auxiliary functions:

$$
z_{cg}(\mathbf{W},k) = \sum_{y=0}^{B} f_{pois}(y|\Lambda_{cg})\exp\left(\sum_{\substack{c'\in\mathcal{N}(c)\\ g'\in\mathcal{N}(g)}}\beta_{gg'}W_{cc'}\arctan(y+k)\arctan(Y_{c'g'})\right) \tag{7}
$$

**Ligand-receptor parameters $\beta_{g_1g_2}$**

The gradient of the Q-function with respect to $\beta_{g_1g_2}$ decomposes into two components:

$$\frac{\partial Q}{\partial \beta_{g_1g_2}} = \sum_{c,g}\left[\left(\sum_{\substack{c'\in N(c)\\ g'\in N(g)}} \frac{\partial \beta_{gg'}}{\partial \beta_{g_1g_2}} e_{cc'}\text{atan}(Y_{cg})\text{atan}(Y_{c'g'})\right) - \frac{\partial \log z_{cg}(\hat{\mathbf{W}}^{(t)})}{\partial \beta_{g_1g_2}}\right]$$

$$= 2\sum_{(c,c')} e_{cc'}\text{atan}(Y_{cg_1})\text{atan}(Y_{c'g_2}) - \sum_{c,g}\frac{1}{z_{cg}(\hat{\mathbf{W}}^{(t)})}\frac{\partial z_{cg}(\hat{\mathbf{W}}^{(t)})}{\partial \beta_{g_1g_2}}$$

The derivative of the normalization factor $z_{cg}$ involves:

$$\frac{\partial z_{cg}(\hat{\mathbf{W}}^{(t)})}{\partial \beta_{g_1g_2}} = \sum_{y=0}^{\infty}\left[\frac{(n_c\lambda_{t(c)g})^y}{y!}\exp(-n_c\lambda_{t(c)g}) \times \exp\left(\sum_{\substack{g'\in N(g)\\ c'\in N(c)}} \beta_{gg'}\hat{W}^{(t)}_{cc'}\text{atan}(y)\text{atan}(Y_{c'g'})\right)\right.$$
$$\left.\times \sum_{\substack{g'\in N(g)\\ c'\in N(c)}} \frac{\partial \beta_{gg'}}{\partial \beta_{g_1g_2}}\hat{W}^{(t)}_{cc'}\text{atan}(y)\text{atan}(Y_{c'g'})\right]$$

Where the partial derivative $\frac{\partial \beta_{gg'}}{\partial \beta_{g_1g_2}} = 1$ if and only if: $(g, g') = (g_1, g_2)$ or $(g_2, g_1)$
We define the auxiliary function $R$:

$$R(g_1,g_2,c,x,\mathbf{W}) = \sum_{y=0}^{B}\left[\frac{(n_c\lambda_{t(c)g_1})^y}{y!}\exp(-n_c\lambda_{t(c)g_1})\right.$$
$$\left.\times \exp\left(\sum_{\substack{g'\in N(g_1)\\ c'\in N(c)}} \beta_{g_1g'}W_{cc'}\text{atan}(y)\text{atan}(Y_{c'g'})\right) \times \left(\sum_{c'\in N(c)} W_{cc'}\text{atan}(y)\text{atan}(Y_{c'g_2})\right)^x\right] \tag{8}$$

Using this notation, the derivatives simplify to:

$$\frac{\partial z_{cg_1}(\hat{\mathbf{W}}^{(t)})}{\partial \beta_{g_1g_2}} = R(g_1, g_2, c, 1, \hat{\mathbf{W}}^{(t)})$$

The complete gradient expression becomes:

$$\frac{\partial Q}{\partial \beta_{g_1g_2}} = 2\sum_{(c,c')} e_{cc'}\text{atan}(Y_{cg_1})\text{atan}(Y_{c'g_2}) - \sum_{c}\left(\frac{R(g_1, g_2, c, 1, \hat{\mathbf{W}}^{(t)})}{z_{cg_1}(\hat{\mathbf{W}})} + \frac{R(g_2, g_1, c, 1, \hat{\mathbf{W}}^{(t)})}{z_{cg_2}(\hat{\mathbf{W}})}\right) \tag{9}$$

The second derivatives:

$$\frac{\partial^2 Q}{\partial \beta^2_{g_1g_2}} = -\sum_{c}\left[\frac{\partial}{\partial \beta_{g_1g_2}}\left(\frac{R(g_1, g_2, c, 1, \hat{\mathbf{W}})}{z_{cg_1}(\hat{\mathbf{W}})}\right) + \frac{\partial}{\partial \beta_{g_1g_2}}\left(\frac{R(g_2, g_1, c, 1, \hat{\mathbf{W}})}{z_{cg_2}(\hat{\mathbf{W}})}\right)\right] \tag{10}$$

Where the individual derivative terms expand as:

$$\frac{\partial}{\partial \beta_{g_1g_2}}\left(\frac{R(g_1, g_2, c, 1, \hat{\mathbf{W}})}{z_{cg_1}(\hat{\mathbf{W}})}\right) = \frac{R(g_1, g_2, c, 2, \hat{\mathbf{W}})z_{cg_1} - R(g_1, g_2, c, 1, \hat{\mathbf{W}})^2}{z^2_{cg_1}}$$

**Cell-type level communication probability $K_{s_1s_2}$**

For cell type pair $(s_1, s_2)$ (where $s_1$ may equal $s_2$), we first define the cell neighborhood set:

$$\mathbf{CNT_{s_1s_2}} := \{(c_1, c_2) : t(c_1) = s_1 \text{ and } t(c_2) = s_2 \text{ or } t(c_2) = s_1 \text{ and } t(c_1) = s_2\}$$

$$
\begin{aligned}
\frac{\partial Q}{\partial K_{s_1 s_2}} &= \sum_{(c_1,c_2)\in \mathbf{CNT}_{s_1 s_2}} (e_{c_1c_2} - \sigma(\eta_{c_1c_2})) \left( \frac{1}{K_{s_1 s_2}} + \frac{1}{1 - K_{s_1 s_2}} \right) \\
\frac{\partial^2 Q}{\partial K_{s_1 s_2}^2} &= \sum_{(c_1,c_2)\in \mathbf{CNT}_{s_1 s_2}} \Bigg[ -\sigma(\eta_{c_1c_2})(1 - \sigma(\eta_{c_1c_2})) \left( \frac{1}{K_{s_1 s_2}} + \frac{1}{1 - K_{s_1 s_2}} \right)^2 \\
&\quad + (e_{c_1c_2} - \sigma(\eta_{c_1c_2})) \left( -\frac{1}{K_{s_1 s_2}^2} + \frac{1}{(1 - K_{s_1 s_2})^2} \right) \Bigg]
\end{aligned} \tag{11}
$$

where $\eta_{c_1c_2} = \gamma_0 + \text{logit}(K_{t(c_1)t(c_2)}) + \gamma_1 d_{c_1c_2}$, $\sigma(x) = (1 + e^{-x})^{-1}$ is the logistic sigmoid function.

**Distance Parameters** $\gamma_0, \gamma_1$

$$
\begin{aligned}
\frac{\partial Q}{\partial \gamma_0} &= \sum_{(c_1,c_2)\in \mathbf{CN}} (e_{c_1c_2} - \sigma(\eta_{c_1c_2})) \\
\frac{\partial^2 Q}{\partial \gamma_0^2} &= -\sum_{(c_1,c_2)\in \mathbf{CN}} \sigma(\eta_{c_1c_2})(1 - \sigma(\eta_{c_1c_2})) \\
\frac{\partial Q}{\partial \gamma_1} &= \sum_{(c_1,c_2)\in \mathbf{CN}} d_{c_1c_2} (e_{c_1c_2} - \sigma(\eta_{c_1c_2})) \\
\frac{\partial^2 Q}{\partial \gamma_1^2} &= -\sum_{(c_1,c_2)\in \mathbf{CN}} d_{c_1c_2}^2 \sigma(\eta_{c_1c_2})(1 - \sigma(\eta_{c_1c_2}))
\end{aligned} \tag{12}
$$

where $\eta_{c_1c_2} = \gamma_0 + \text{logit}(K_{t(c_1)t(c_2)}) + \gamma_1 d_{c_1c_2}$,$\sigma(x) = (1 + e^{-x})^{-1}$ is the logistic sigmoid function.

**Convergence Criterion**

Terminate when:

$$
\max_{\theta\in\Theta} |\theta^{(t)} - \theta^{(t-1)}| < \epsilon
$$

where $\Theta = \{\lambda, \beta, \mathbf{K}, \gamma_0, \gamma_1\}$.

Finally, we summarize the algorithm as follows:

**Algorithm 1:** ICM-EM algorithm for IC3 model

**Data:** count matrix $\mathbf{Y}$, cell type information $\mathbf{T}$, ligand-receptor database $\mathbf{LR}$, communication threshold distance $d_0$, cell location matrix $\mathbf{L}$

**Result:** Cell type communication probability matrix $\mathbf{K}$, the posterior probability of cell type pairs $e$, Poisson distribution parameter $\lambda$,communication strength of ligand receptor gene pair $\beta$.

Compute cell distance matrix $\mathbf{D}$ by cell location matrix $\mathbf{L}$.<br>
Generate $\mathbf{CN}$,$\mathbf{CNT_{s_1 s_2}}$ by $\mathbf{D}$, $d_0$ and $\mathbf{T}$.<br>
Generate $N(c)$,$N(g)$ for each cell c and gene g by $\mathbf{CN}$ and $\mathbf{J}$.<br>
Initialize all parameters $\Theta^0 = (\lambda^{(0)}, \beta^{(0)}, \mathbf{K}^{(0)}, \gamma_0^{(0)}\gamma_1^{(0)}$ by initialization step.<br>
$itr \leftarrow 1$;<br>
**while** $\Theta^{change} > \varepsilon$ **do**<br>
  **E step**<br>
  **for** $(c_1, c_2) \in \mathbf{CP}$ **do**<br>
    Calculate $b_1(c_1, c_2), b_2(c_1, c_2)$ from eq3,eq4;<br>
    Update $e_{c_1 c_2}^{(itr)}$ from eq1;<br>
    Update $\hat{W}_{c_1 c_2}^{(itr)}$ from eq5;<br>
  **M step**<br>
  **for** $g \in \mathbf{Gene}; t \in \mathbf{Type}$ **do**<br>
    Calculate $z_{cg}^1(\hat{\mathbf{W}}^{(t)})$, $z_{cg}^2(\hat{\mathbf{W}}^{(t)})$ from eq7;<br>
    Calculate $\frac{\partial Q}{\partial \lambda_{kg}}, \frac{\partial Q^2}{\partial^2 \lambda_{kg}}$ from eq6;<br>
    Update $\lambda_{kg}$ by $\lambda_{kg}^{(itr)} = \lambda_{kg}^{(itr-1)} - \frac{\frac{\partial Q}{\partial \lambda_{kg}}}{\frac{\partial Q^2}{\partial^2 \lambda_{kg}}}$.<br>
  **for** $s_1 \in \mathbf{Type}; s_2 \in \mathbf{Type}$ **do**<br>
    **for** $(g_1, g_2) \in \mathbf{LR}$ **do**<br>
      **for** $c \in \{c : t(c) = s_1\}$ **do**<br>
        Calculate $R(g_1, g_2, c, s_2, 1, \hat{\mathbf{W}}^{(t)})$, $R(g_1, g_2, c, s_2, 2, \hat{\mathbf{W}}^{(t)})$ from eq8;<br>
      **for** $c \in \{c : t(c) = s_2\}$ **do**<br>
        Calculate $R(g_1, g_2, c, s_1, 1, \hat{\mathbf{W}}^{(t)})$, $R(g_1, g_2, c, s_1, 2, \hat{\mathbf{W}}^{(t)})$ from eq8;<br>
      Calculate $\frac{\partial Q}{\partial \beta_{g_1 g_2 c_1 c_2}}$ from (28);<br>
      Calculate $\frac{\partial Q^2}{\partial^2 \beta_{g_1 g_2 c_1 c_2}}$ from eq9;<br>
      Update $\beta_{g_1 g_2 c_1 c_2}$ by $\beta_{g_1 g_2 c_1 c_2}^{(itr)} = \beta_{g_1 g_2 c_1 c_2}^{(itr-1)} - \frac{\frac{\partial Q}{\partial \beta_{g_1 g_2 c_1 c_2}}}{\frac{\partial Q^2}{\partial^2 \beta_{g_1 g_2 c_1 c_2}}}$. from eq10<br>
  **for** $s_1 \in \mathbf{Type}; s_2 \in \mathbf{Type}$ **do**<br>
    Calculate $\frac{\partial Q}{\partial K_{s_1 s_2}}, \frac{\partial^2 Q}{\partial K_{s_1 s_2}^2}$ from eq11;<br>
    Update $K_{s_1 s_2}$ by $K_{s_1 s_2}^{(itr)} = K_{s_1 s_2}^{(itr-1)} - \frac{\frac{\partial Q}{\partial K_{s_1 s_2}}}{\frac{\partial Q^2}{\partial^2 K_{s_1 s_2}}}$.<br>
  Calculate $\frac{\partial Q}{\partial \gamma_0}, \frac{\partial^2 Q}{\partial \gamma_0^2}, \frac{\partial Q}{\partial \gamma_1}, \frac{\partial^2 Q}{\partial \gamma_1^2}$ from eq12;<br>
  Update $\gamma_0$ by $\gamma_0^{(itr)} = \gamma_0^{(itr-1)} - \frac{\frac{\partial Q}{\partial \gamma_0}}{\frac{\partial Q^2}{\partial^2 \gamma_0}}$. Update $\gamma_0$ by $\gamma_0^{(itr)} = \gamma_1^{(itr-1)} - \frac{\frac{\partial Q}{\partial \gamma_1}}{\frac{\partial Q^2}{\partial^2 \gamma_1}}$;<br>
  Calculate $\Theta^{\text{change}}$ ;<br>
  $itr \leftarrow itr + 1$;

### 2 Supplementary Note 2: Simulation studies

#### 2.1 Sparsity level

We simulated spatial transcriptomic data based on the seqFISH+ olfactory bulb (OB) dataset [3]. We used the spatial positions of the 252 cells in dataset and simulated cell types and the expression levels of ligands and receptors of these cells. The simulated data divided the 252 cells into 3 main cell types and 2 rare cell types. We assume that the proportion of main cell types in all cells is $p_{\text{main}}$, and the proportion of rare cell types in all cells was $p_{\text{rare}}$. The sparsity of the simulated data is defined as follows.

$$\textbf{Sparsity} = \frac{p_{\text{rare}}}{p_{\text{main}}}$$

#### 2.2 Data generation details

We constructed datasets at 10 sparsity levels, where the sparsity parameter $\delta$ was set to 0.1, 0.2, 0.3,..., 0.9 and 1, respectively. For a given sparsity $\delta$, the cell type proportions of the five cell populations followed a ratio of 1:1:1:$\delta$:$\delta$. Cell type labels were then randomly sampled according to these predefined proportions. For each sparsity level, we repeated the simulation 100 times.

For the probability of communication at the cell type level $\mathbf{K}$, each element $\mathbf{K}_{ij}$ is independently sampled from a Bernoulli distribution with probability 0.3.

In constructing the cell-level communication network, we assign $\mathbf{W} = 1$ between adjacent cells when they belong to cell types that are known to engage in interaction.

The ligand-receptor parameters $\beta$ are drawn from a zero-inflated exponential distribution:

$$\beta \sim 0.7\delta_0 + 0.3\text{Exp}(1)$$

where $\delta_0$ denotes a point mass at zero and Exp(1) represents the exponential distribution with rate parameter 1.

We used the mean expression values of genes in the OB data to represent the Poisson parameters of these genes in the simulated data.

The simulation of single-cell transcriptomic data proceeds through an Gibbs sampling framework. After initialization, the expression value of each gene is sampled from the conditional distribution determined by the graphical model. This process repeats sequentially across all genes and cells for 100 iterations, with the final iteration yielding the simulated transcriptomic data.

#### 2.3 Toy model details

We constructed a toy model consisting of three cells, where the middle cell secretes ligands, and the two side cells secrete receptors. The expression level of the ligands secreted by the middle cell was fixed at 3, while the expression levels of receptors secreted by the side cells could vary from 0 to 10, resulting in a total of 11 × 11 possible combinations. Our goal is to infer the values of $W_1$ and $W_2$ under these 121 conditions. Since this toy model involves only two latent variables, the ICM-EM algorithm is unnecessary for inference. Instead, we can directly use the EM algorithm. In the E-step, we enumerate all four possible combinations of values for $W_1$ and $W_2$, compute their corresponding expectations, and thus obtain the Q function.

### 3 Supplementary Note 3: The scan statistic and permutation test

We use the scan statistic [4] to identify the communication hotspots. A slice in the MERFISH dataset of hypothalamic preoptic region [5] is a square of 1800 µm by 1800 µm. We scan for hotspots with squares, the side lengths varying between 40 µm and 540 µm with a common difference of 25 µm, to identify the square with optimal likelihood ratio.

For a particular slice and a pair of cell types (A, B) under study, the optimal score is $s^{AB}$. Next we perform permutation test to obtain the null distribution of $s^{AB}$. Let $CN^{\text{AB}}$ be the set of cell pairs of type A and B with distance less than $d_0$ in the slice. Let $n^{\text{AB}}$ be the number of cell pairs of type A and B with posterior communication probability greater than 0.5. In each permutation, we randomly extract $n^{\text{AB}}$ pairs of cells from $CN^{\text{AB}}$. These pairs are considered to have cell-cell communications , while the other pairs are considered not to have cell-cell communications. For this permutated network, we do the same search to obtain the optimal likelihood ratio (as above). The optimal score for the $i$-th permuation is denoted as $s^{\text{AB}}_{(i)}$. The permuation is repeated for $1,000$ times. The empirical P value of $s^{\text{AB}}$ is calculated as:

$$\text{P value} = \frac{|\{i : |s^{\text{AB}}_{(i)} \geq s^{\text{AB}}\}|}{1000}$$

If the p-value is 0, we can assume that the null distribution of $s^{\mathrm{AB}}$ follows a normal distribution; we get the p-value as follows:

$$\mu_s = \frac{\sum_{i=1}^{N} s_{(i)}^{\mathrm{AB}}}{N}; \sigma_s^2 = \frac{\sum_{i=1}^{N} (s_{(i)}^{\mathrm{AB}} - \mu_s)^2}{N-1}; \quad \text{P value} = P(X > \frac{s^{\mathrm{AB}} - \mu_s}{\sigma_s^2}), X \sim Normal(0,1)$$

### 4 Supplementary Tables

| Type 1 | Type 2 | Pubmed ID |
|---|---|---|
| Astrocyte | Mitral/Tufted cell | 10570454 |
| Olfactory ensheathing cell | Endothelial cell | 15319002 |
| Olfactory ensheathing cell | Astrocyte | 17611269 |
| Olfactory ensheathing cell | Olfactory ensheathing cell | 20143249 |
| Olfactory ensheathing cell | Astrocyte | 20713050 |
| Neuroblast | Endothelial cell | 24322013 |
| Neuroblast | Microglia | 25076873 |
| Endothelial cell | Endothelial cell | 25283993 |
| Mitral/Tufted cell | Interneuron | 25762678 |
| Interneuron | Astrocyte | 26586821 |
| Neuroblast | Astrocyte | 28258169 |
| Neuroblast | Neuroblast | 28258169 |
| Astrocyte | Astrocyte | 28258169 |
| Interneuron | Olfactory ensheathing cell | 28674010 |
| Astrocyte | Microglia | 30282022 |
| Astrocyte | Oligodendrocyte | 30692691 |
| Endothelial cell | Microglia | 31221990 |
| Mitral/Tufted cell | Mitral/Tufted cell | 33580583 |
| Interneuron | Interneuron | 34843626 |
| Interneuron | Microglia | 34919273 |
| Neuroblast | Oligodendrocyte | 36185360 |
| Neuroblast | Mitral/Tufted cell | 20397619 |
| Astrocyte | Endothelial cell | 16371949 |
| Interneuron | Oligodendrocyte | 38612430 |

**Table 1: The benchmark dataset of cell-cell communications for the Olfactory bulb region in mouse brain. The first two columns indicate the two cell types, and the third column provides the PubMed ID of the supporting literature.**

| Type 1 | Type 2 | PubMed ID |
|---|---|---|
| Neuroblast | Astrocyte | 10377448 |
| Neural progenitor cell | Microglia | 30177874 |
| Ependymal cell | Neural progenitor cell | 30177874 |
| Ependymal cell | Neuroblast | 30177874 |
| Ependymal cell | Neural stem cell | 30177874 |
| Interneuron | Ependymal cell | 35450289 |
| Neural stem cell | Neural progenitor cell | 20844536 |
| Neuroblast | Neuroblast | 17883412 |
| Neural stem cell | Endothelial cell | 21253820 |
| Interneuron | Astrocyte | 23619383 |
| Oligodenrocyte progenitor cell | Neuroblast | 23671797 |
| Microglia | Neural stem cell | 25245208 |
| Neural progenitor cell | Astrocyte | 26528139 |
| Neural progenitor cell | Microglia | 26528139 |
| Neural stem cell | Endothelial cell | 28694515 |
| Interneuron | Interneuron | 30177874 |
| Neural progenitor cell | Neural progenitor cell | 30177874 |
| Neural stem cell | Interneuron | 30777863 |
| Neural stem cell | Choroid plexus | 30777863 |
| Neural stem cell | Neural stem cell | 30777863 |
| Neuroblast | Choroid plexus | 31064838 |
| Endothelial cell | Neural progenitor cell | 32502570 |
| Neuroblast | Oligodendrocyte | 36185360 |
| Astrocyte | Oligodendrocyte | 37291151 |
| Choroid plexus | Astrocyte | 26174708 |
| Choroid plexus | Interneuron | 26174708 |
| Choroid plexus | Choroid plexus | 26174708 |
| Ependymal cell | Ependymal cell | 36415662 |
| Oligodendrocyte | Oligodenrocyte progenitor cell | 31770564 |

**Table 2: TThe benchmark dataset of cell-cell communications for the subventricular zone in mouse brain. The first two columns indicate the two cell types, and the third column provides the PubMed ID of the supporting literature.**

| Type 1 | Type 2 | PubMed ID |
|---|---|---|
| Astrocyte | Astrocyte | 19807846 |
| Astrocyte | Endothelial cell | 11999726 |
| Astrocyte | Oligodendrocyte | 11999726 |
| Astrocyte | Microglia | 29744298 |
| Endothelial cell | Endothelial cell | 11999726 |
| Endothelial | Pericytes | 32485292 |
| Excitatory | Excitatory | 14579004 |
| Excitatory | Inhibitory | 14579004 |
| Inhibitory | Inhibitory | 14579004 |
| Microglia | Oligodendrocytes | 36199097 |
| Microglia | Inhibitory | 34233165 |
| Astrocyte | Excitatory | 37563296 |
| Astrocyte | Inhibitory | 37563296 |
| Astrocyte | Endothelial | 29411111 |

**Table 3: The benchmark dataset of cell-cell communications for the hypothalamic preoptic region in mouse brain. The first two columns indicate the two cell types, and the third column provides the PubMed ID of the supporting literature.**

#### 5 Supplementary Figures

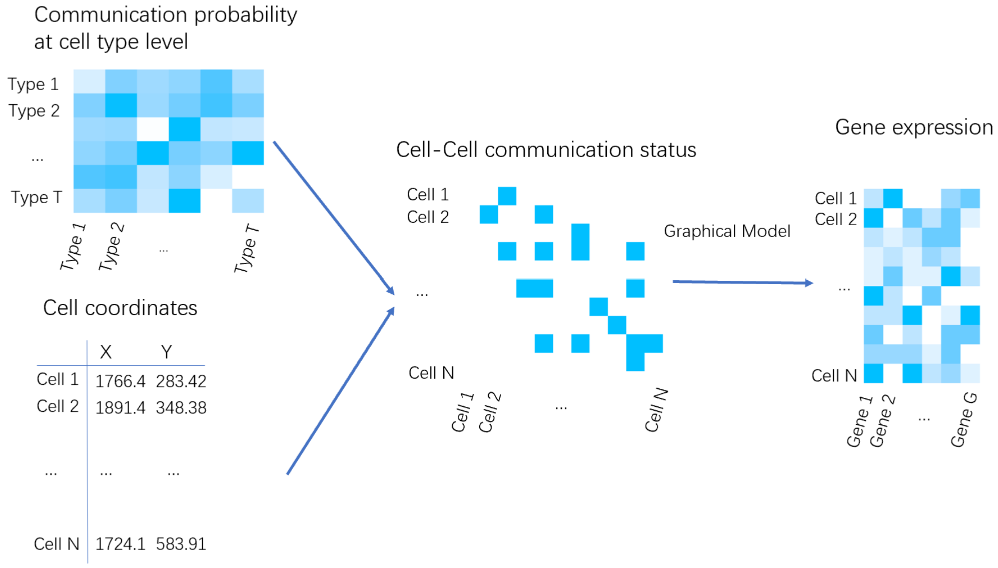

**Figure S1: Overview of IC3 model. IC3 is a hierarchical graphical model. For the first layer, the cell-type level communication probability and the distance matrix determines the distribution of the communication status. At the second layer, the latent communication network determines the distribution of the expression levels of ligands and receptors.**

(a) (b)

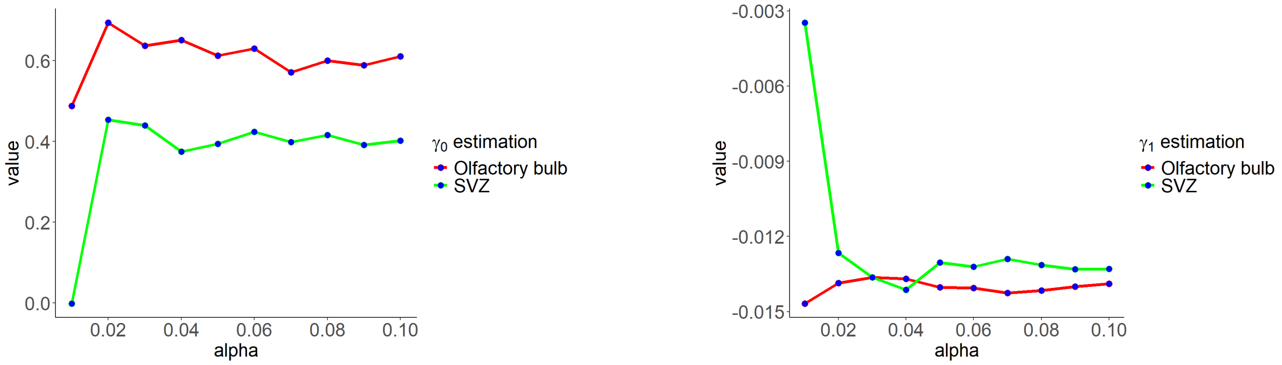

**Figure S2: $\gamma$ Sensitivity analysis of the choice of $d_0$. Suppose $Dv$ is the numerical vector of distances between any two cells in the dataset. $d_0$ is the $\alpha$ upper quantile of $Dv$. (a),(b): The estimation of the parameters $\gamma_0$ and $\gamma_1$ with varying levels of $\alpha$.**

(a) (b)

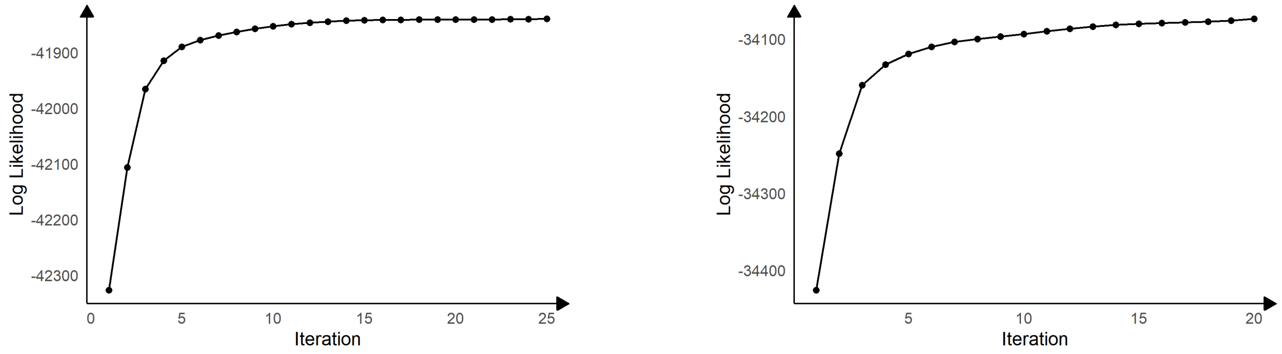

**Figure S3: Convergence of the value of Q function in the ICM-EM algorithm (a): SVZ (b):Olfactory bulb.**

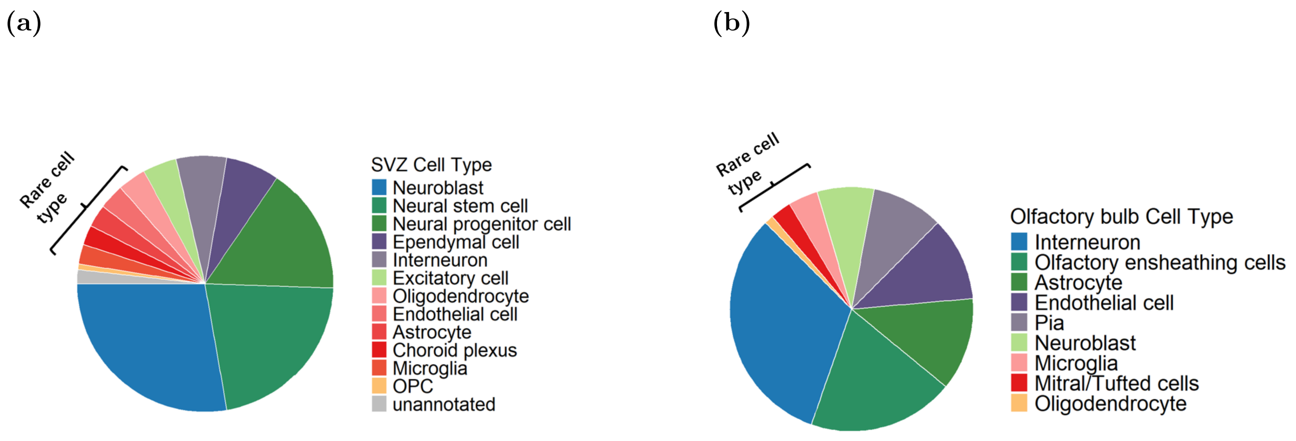

**Figure S4: The proportion of different cell types in the example slice in SVZ (a) and OB (b) regions. The rare cell types (fewer than 10 cells) are represented in warm colors.**

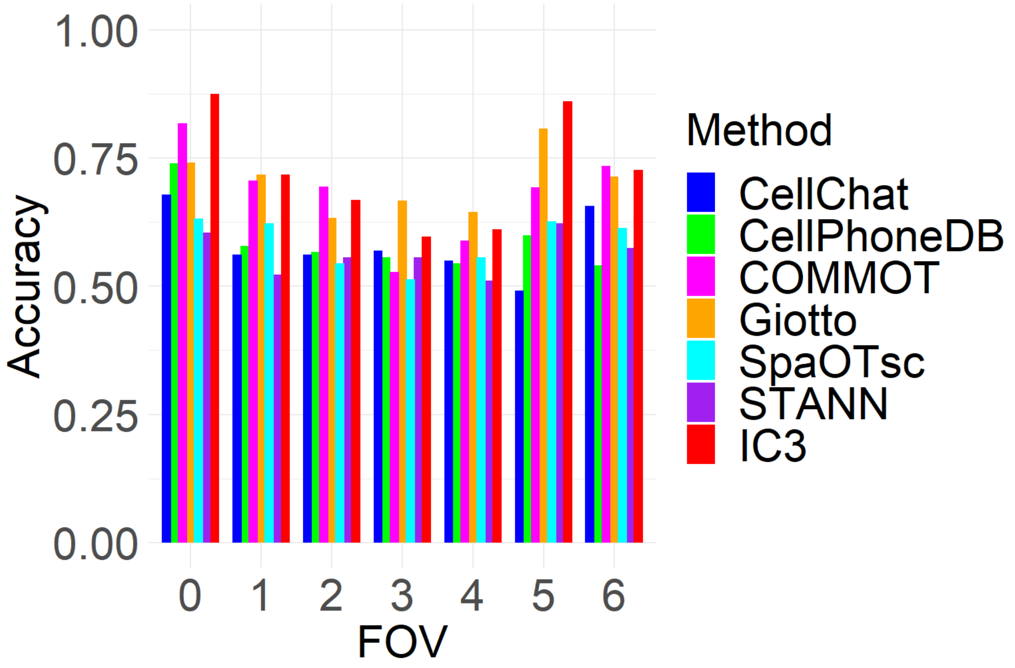

**Figure S5: Bar chart of AUC results for different methods across 7 FOVs for the OB region.**

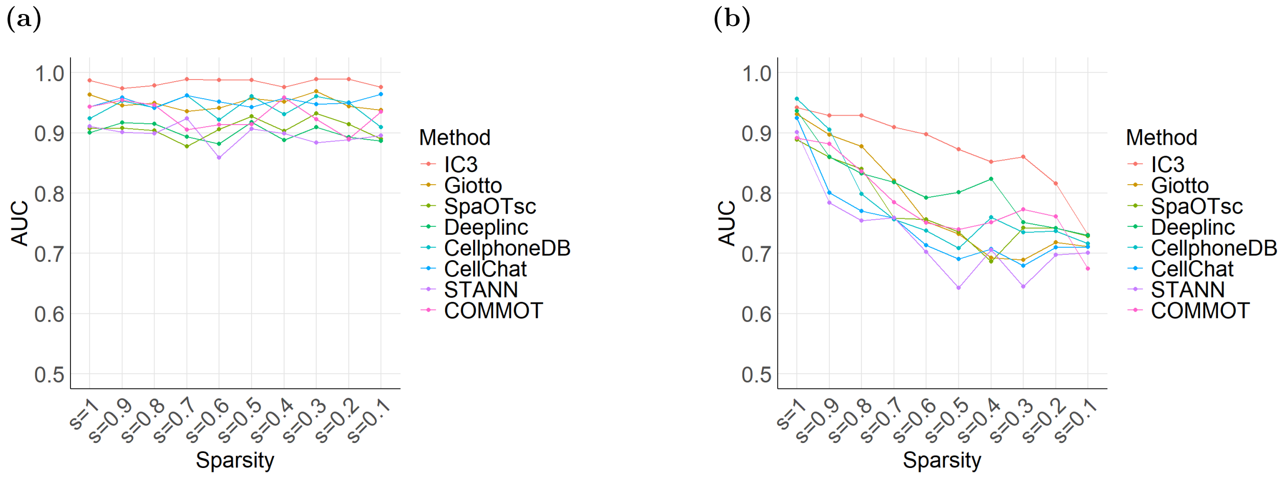

**Figure S6: Comparison of AUC values at varying sparsity levels for the IC3 method and the other seven methods for non-rare cell types (a) and rare cell types (b).**

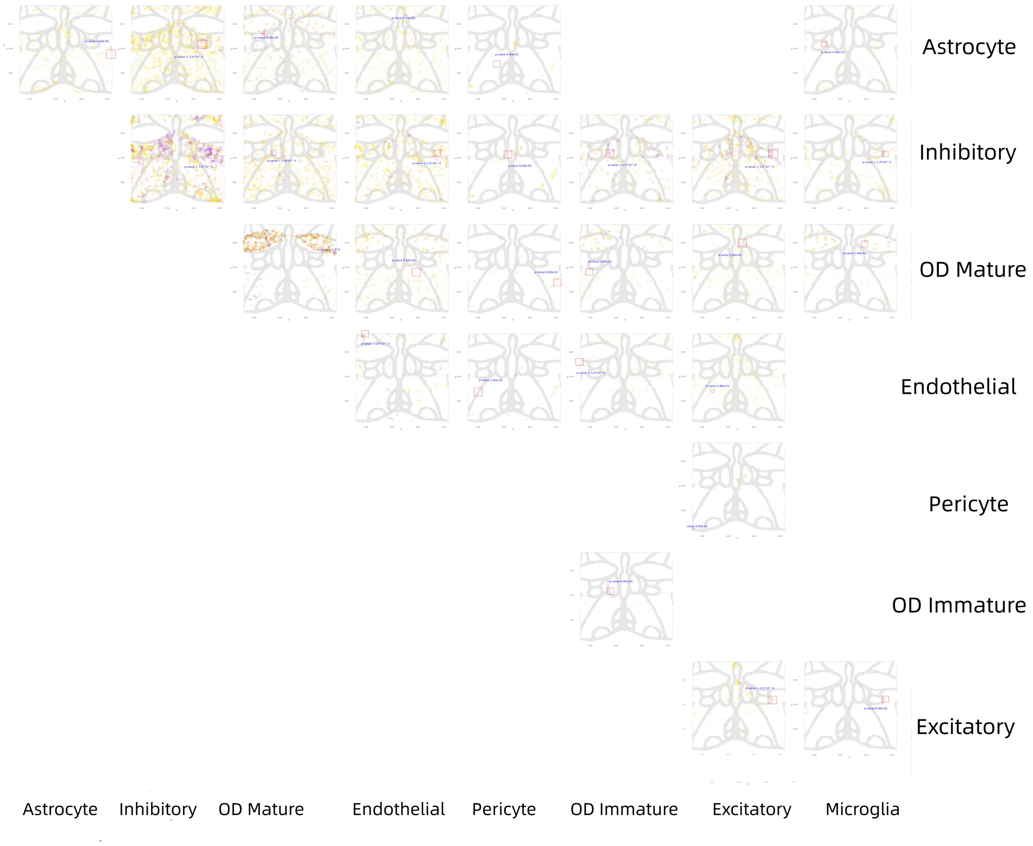

**Figure S7: The communication network of Slice 1 in the MERFISH dataset of mouse brain. Purple dots indicate a cell-cell communications, and the location of the dot is the middle point of the two cells. Red rectangle highlight significant communication hotspots.**

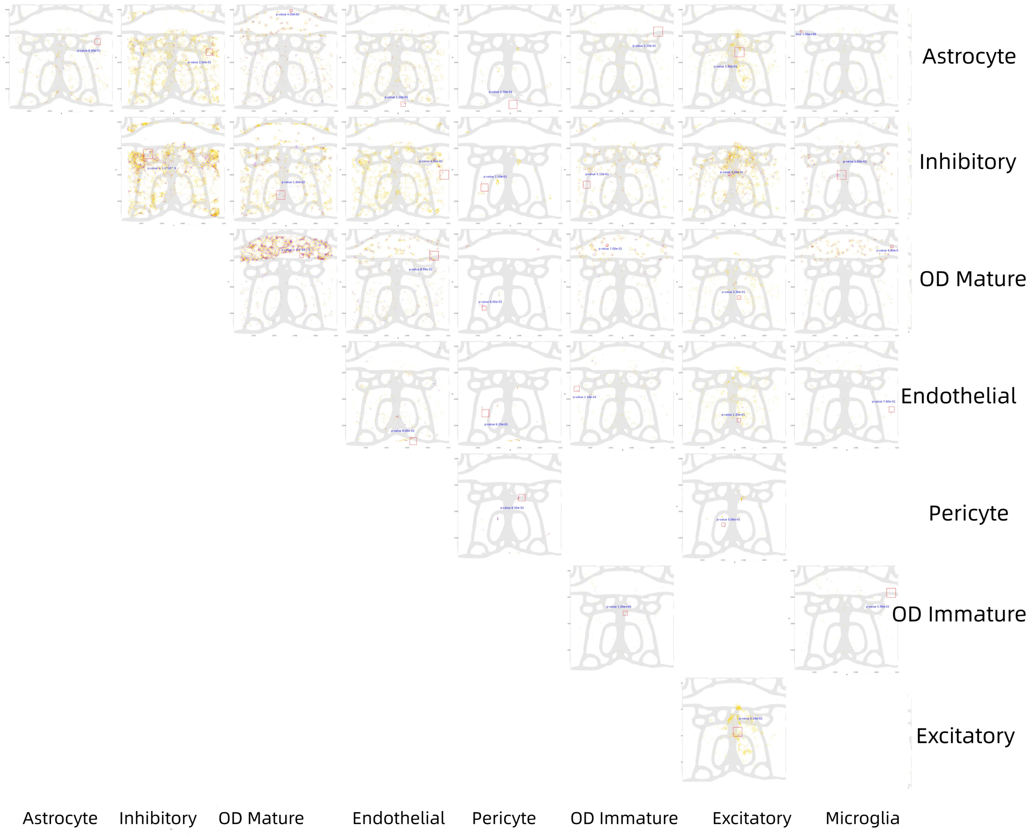

**Figure S8: The communication network of Slice 2 in the MERFISH dataset of mouse brain. Purple dots indicate a cell-cell communications, and the location of the dot is the middle point of the two cells. Red rectangle highlight significant communication hotspots.**

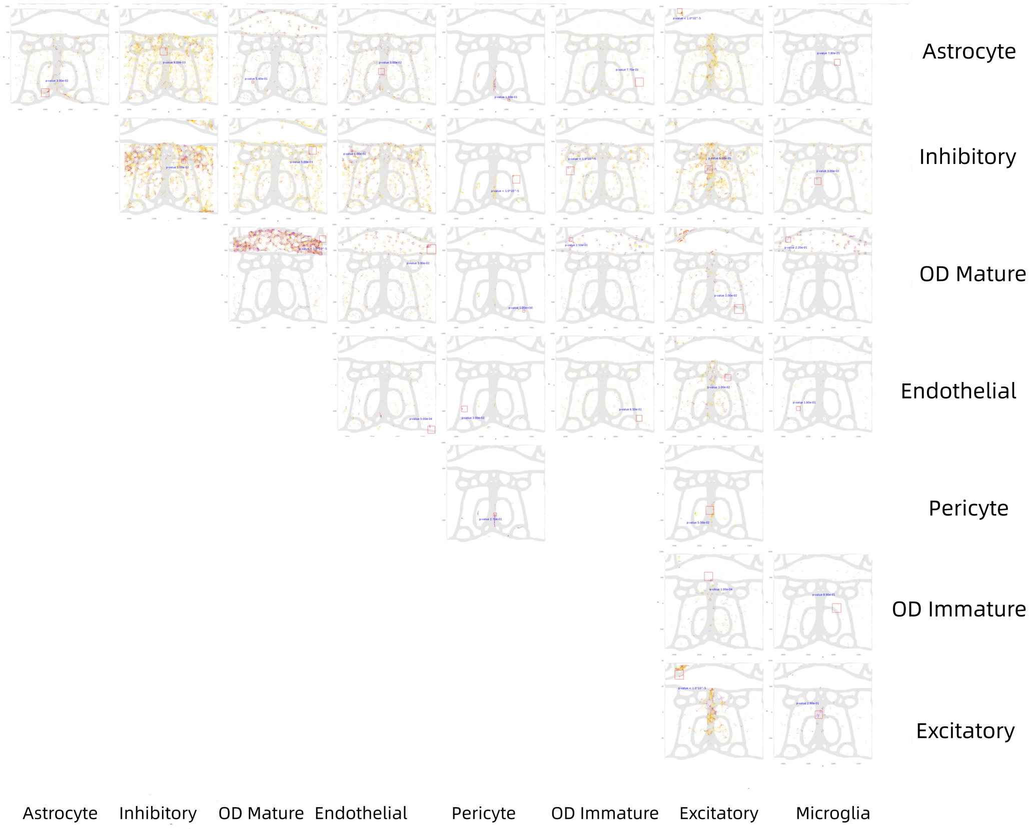

**Figure S9: The communication network of Slice 3 in the MERFISH dataset of mouse brain. Purple dots indicate a cell-cell communications, and the location of the dot is the middle point of the two cells. Red rectangle highlight significant communication hotspots.**

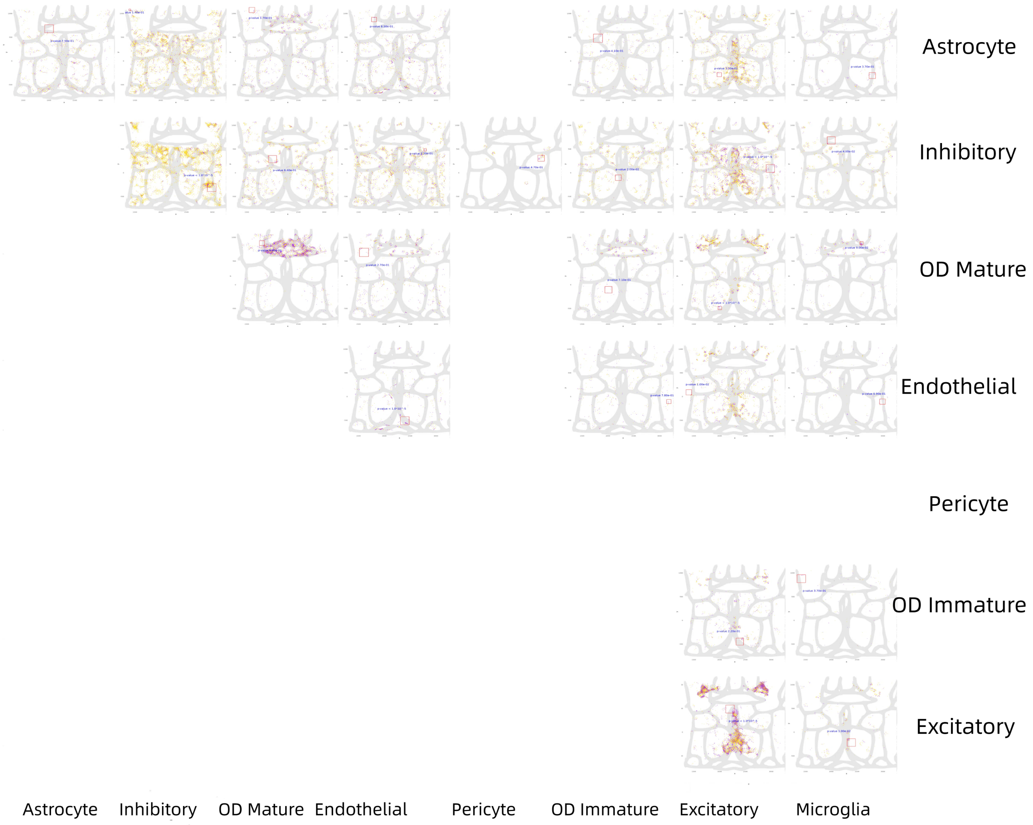

**Figure S10: The communication network of Slice 4 in the MERFISH dataset of mouse brain. Purple dots indicate a cell-cell communications, and the location of the dot is the middle point of the two cells. Red rectangle highlight significant communication hotspots.**

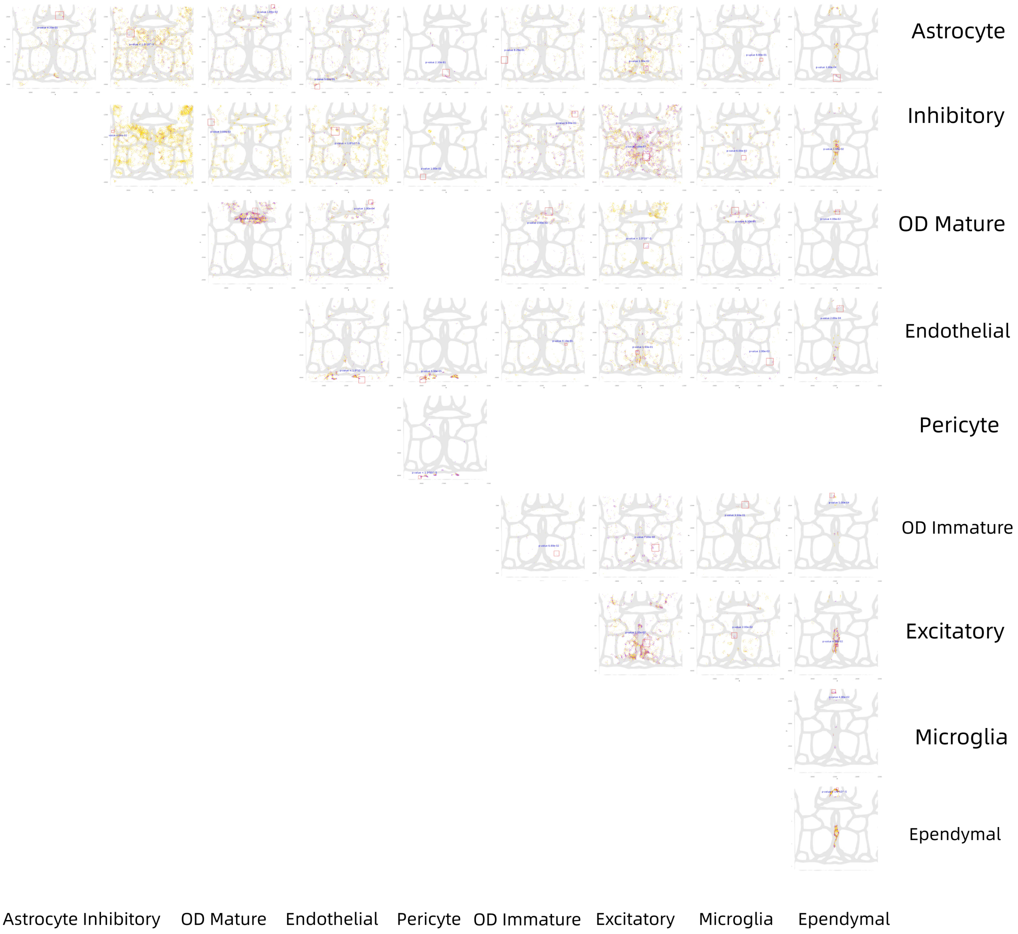

**Figure S11: The communication network of Slice 5 in the MERFISH dataset of mouse brain. Purple dots indicate a cell-cell communications, and the location of the dot is the middle point of the two cells. Red rectangle highlight significant communication hotspots.**

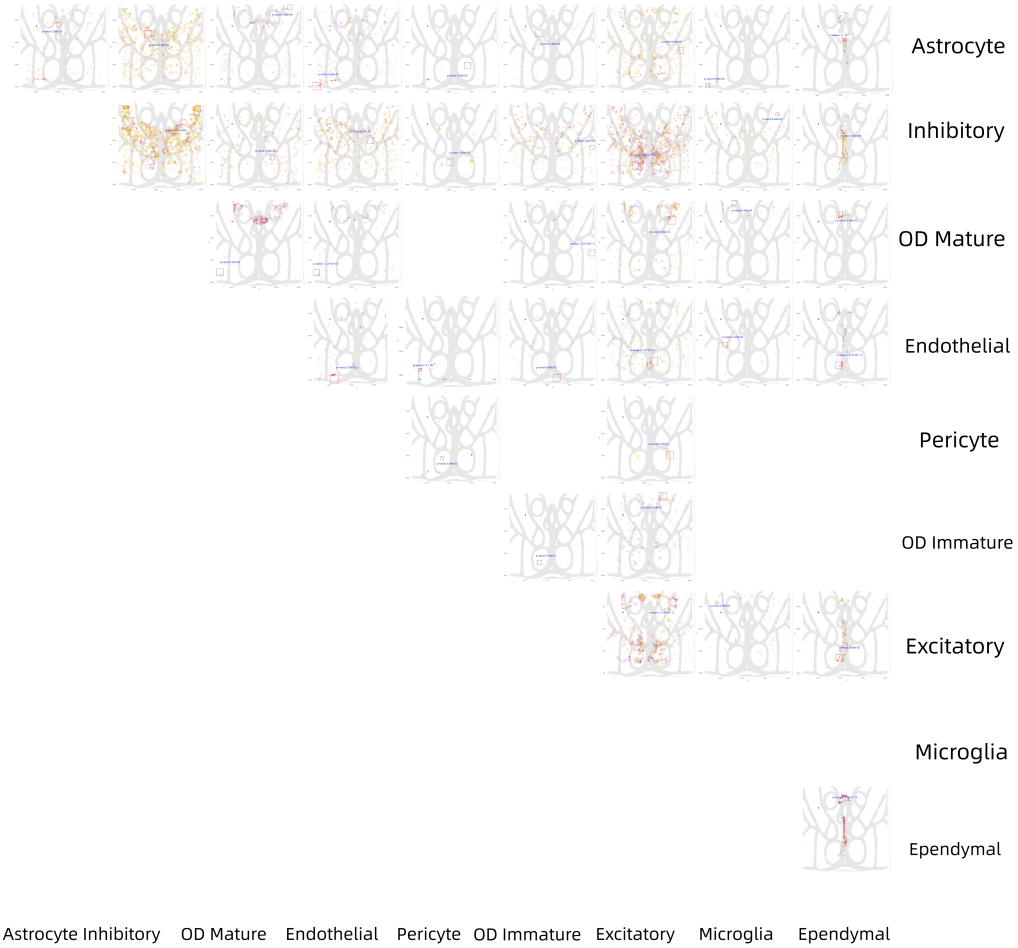

**Figure S12: The communication network of Slice 6 in the MERFISH dataset of mouse brain. Purple dots indicate a cell-cell communications, and the location of the dot is the middle point of the two cells. Red rectangle highlight significant communication hotspots.**

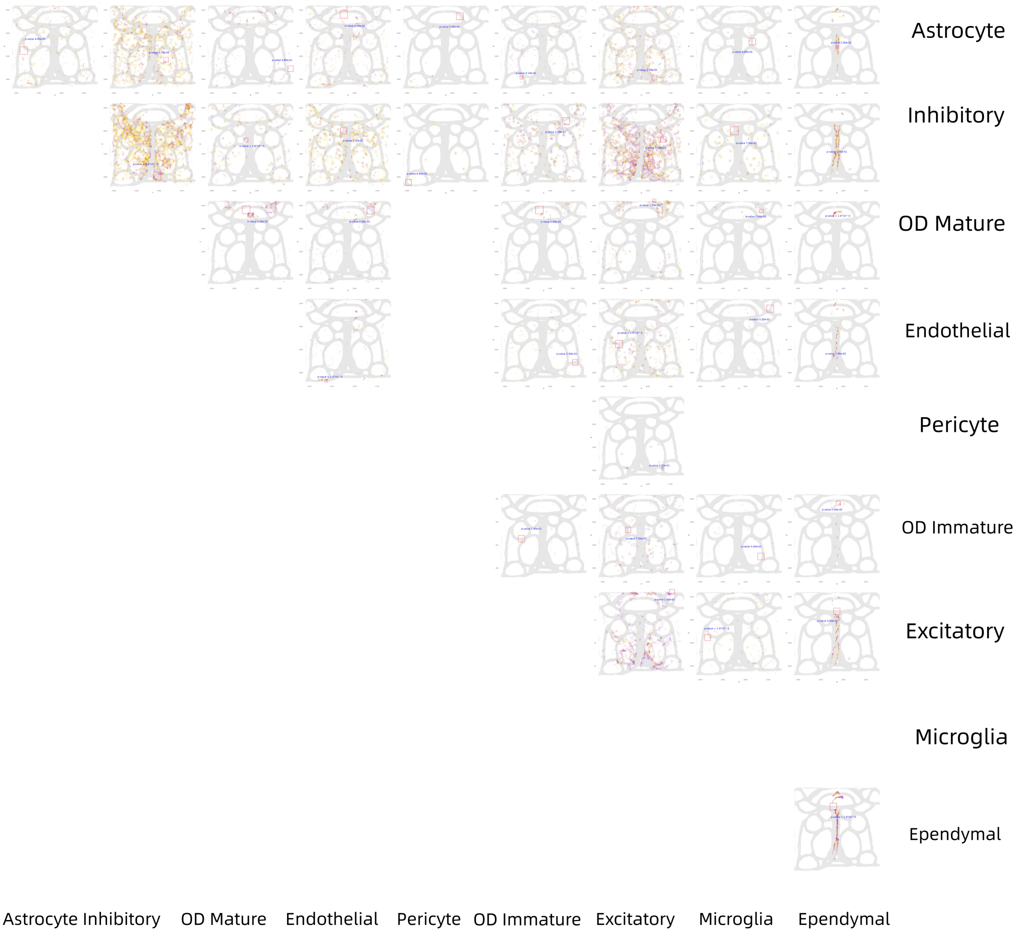

**Figure S13: The communication network of Slice 7 in the MERFISH dataset of mouse brain. Purple dots indicate a cell-cell communications, and the location of the dot is the middle point of the two cells. Red rectangle highlight significant communication hotspots.**

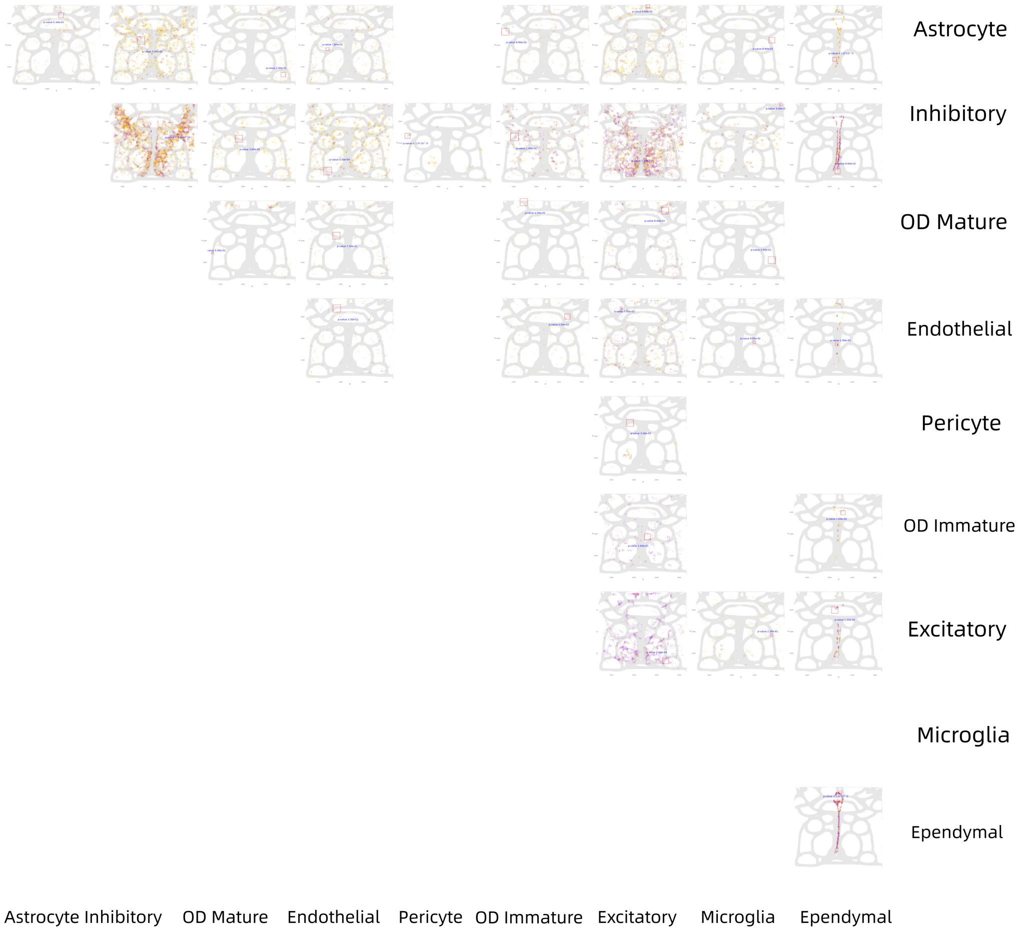

**Figure S14: The communication network of Slice 8 in the MERFISH dataset of mouse brain. Purple dots indicate a cell-cell communications, and the location of the dot is the middle point of the two cells. Red rectangle highlight significant communication hotspots.**

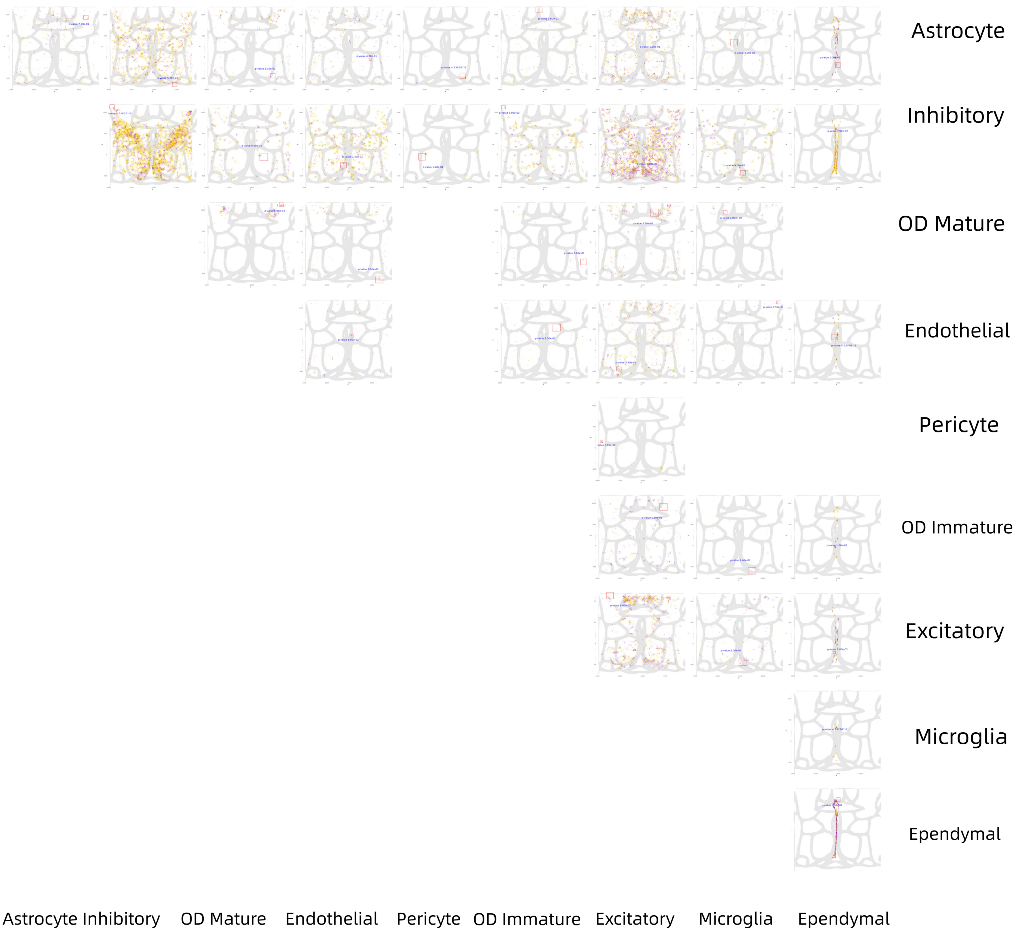

**Figure S15: The communication network of Slice 9 in the MERFISH dataset of mouse brain. Purple dots indicate a cell-cell communications, and the location of the dot is the middle point of the two cells. Red rectangle highlight significant communication hotspots.**

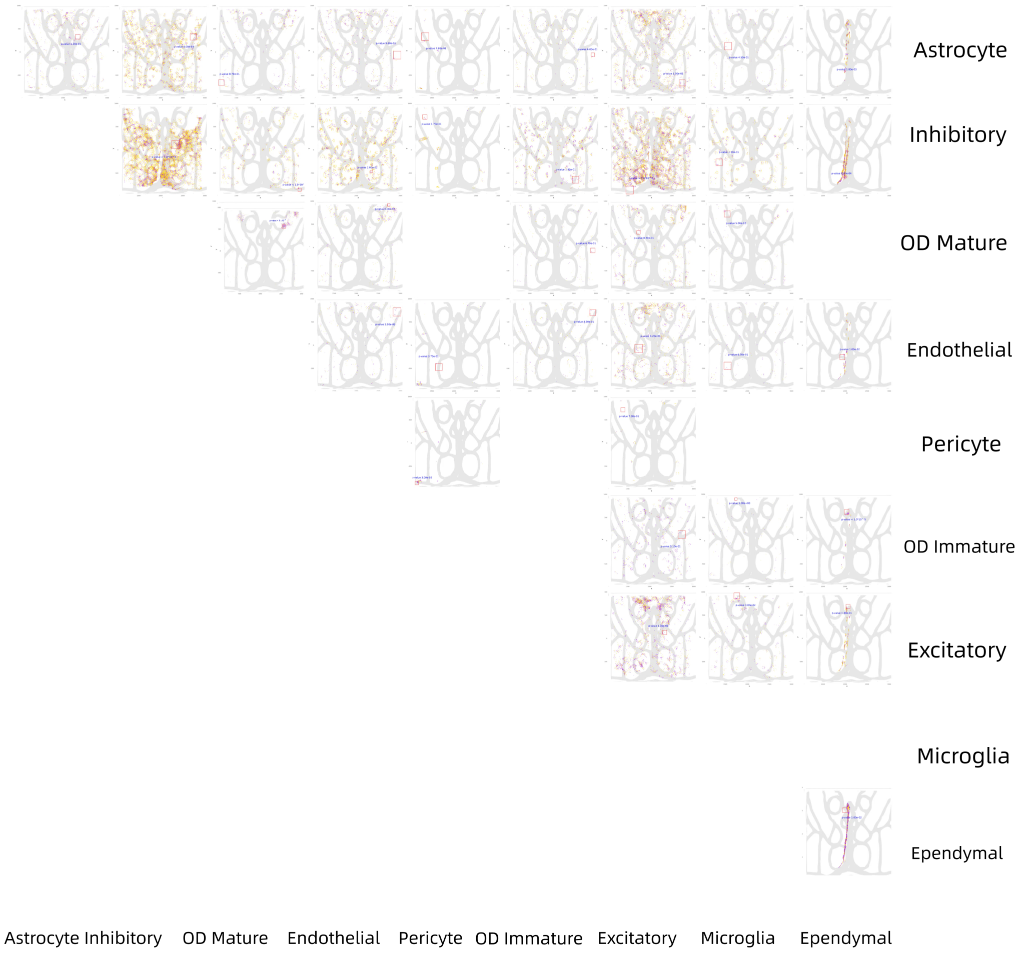

**Figure S16: The communication network of Slice 10 in the MERFISH dataset of mouse brain. Purple dots indicate a cell-cell communications, and the location of the dot is the middle point of the two cells. Red rectangle highlight significant communication hotspots.**

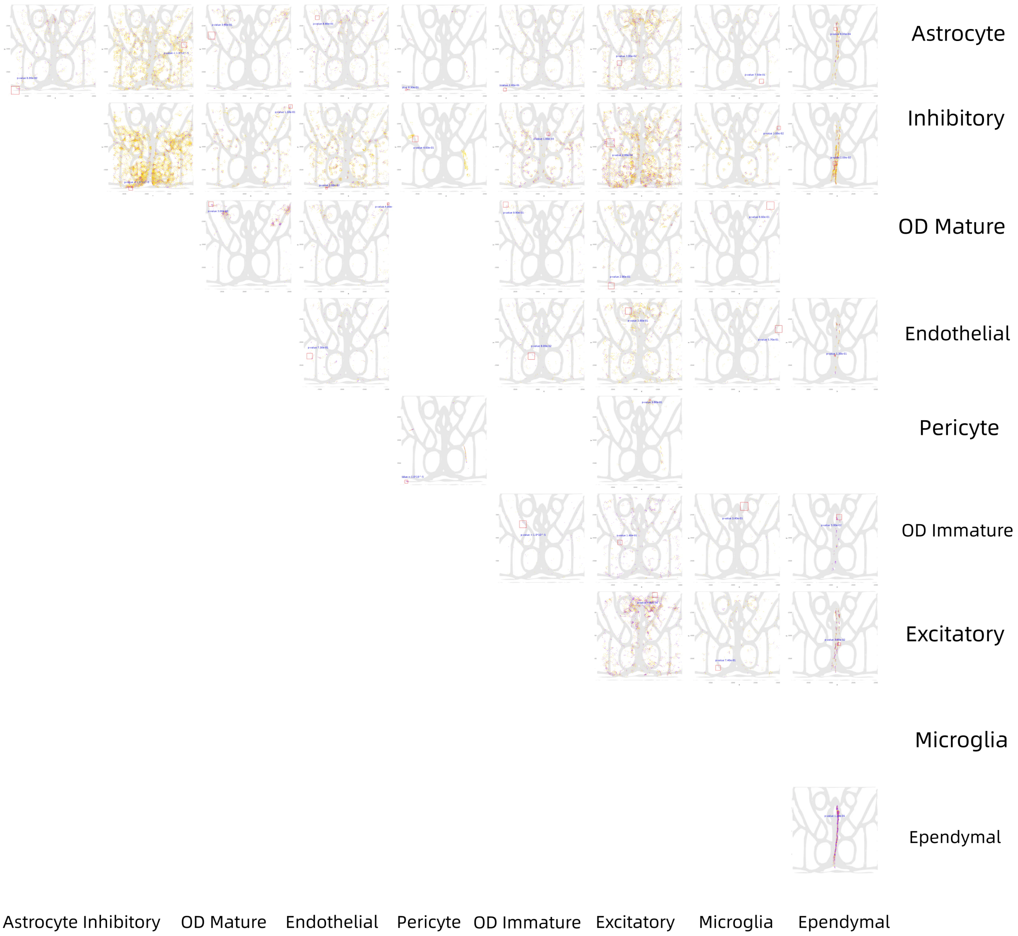

**Figure S17: The communication network of Slice 11 in the MERFISH dataset of mouse brain. Purple dots indicate a cell-cell communications, and the location of the dot is the middle point of the two cells. Red rectangle highlight significant communication hotspots.**

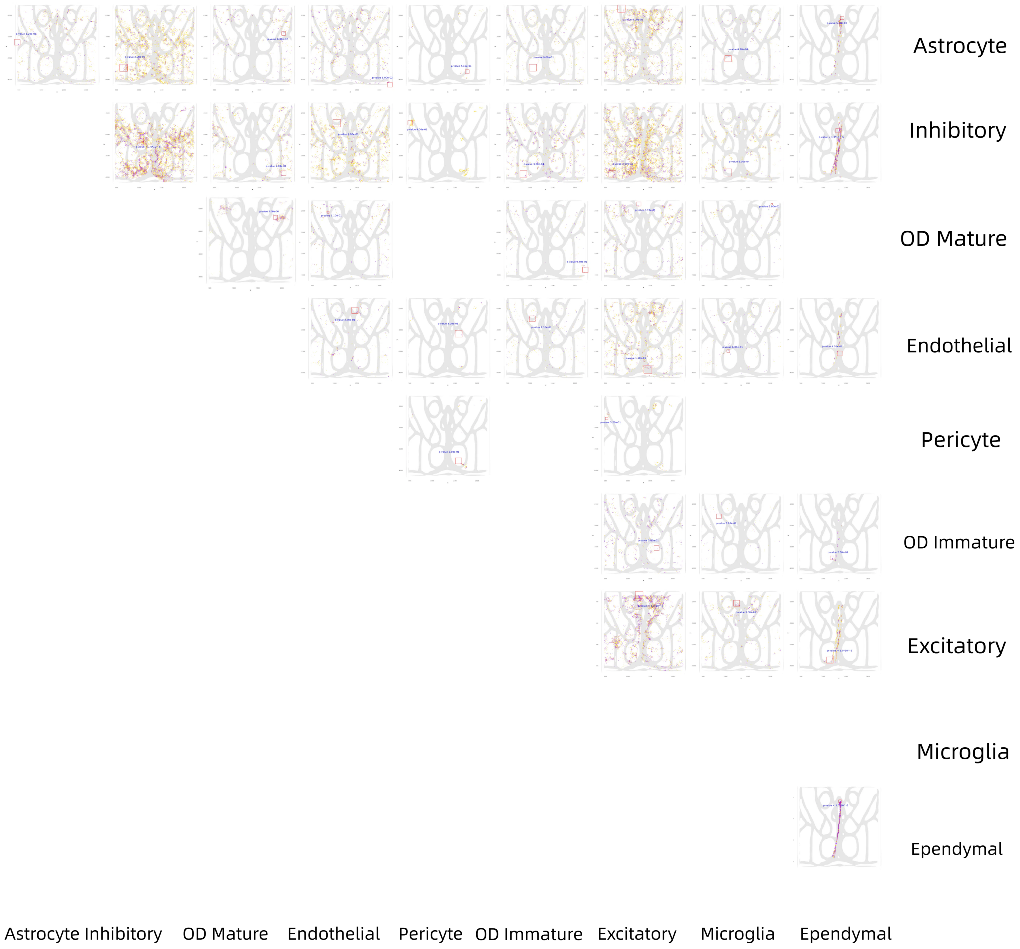

**Figure S18: The communication network of Slice 12 in the MERFISH dataset of mouse brain. Purple dots indicate a cell-cell communications, and the location of the dot is the middle point of the two cells. Red rectangle highlight significant communication hotspots.**

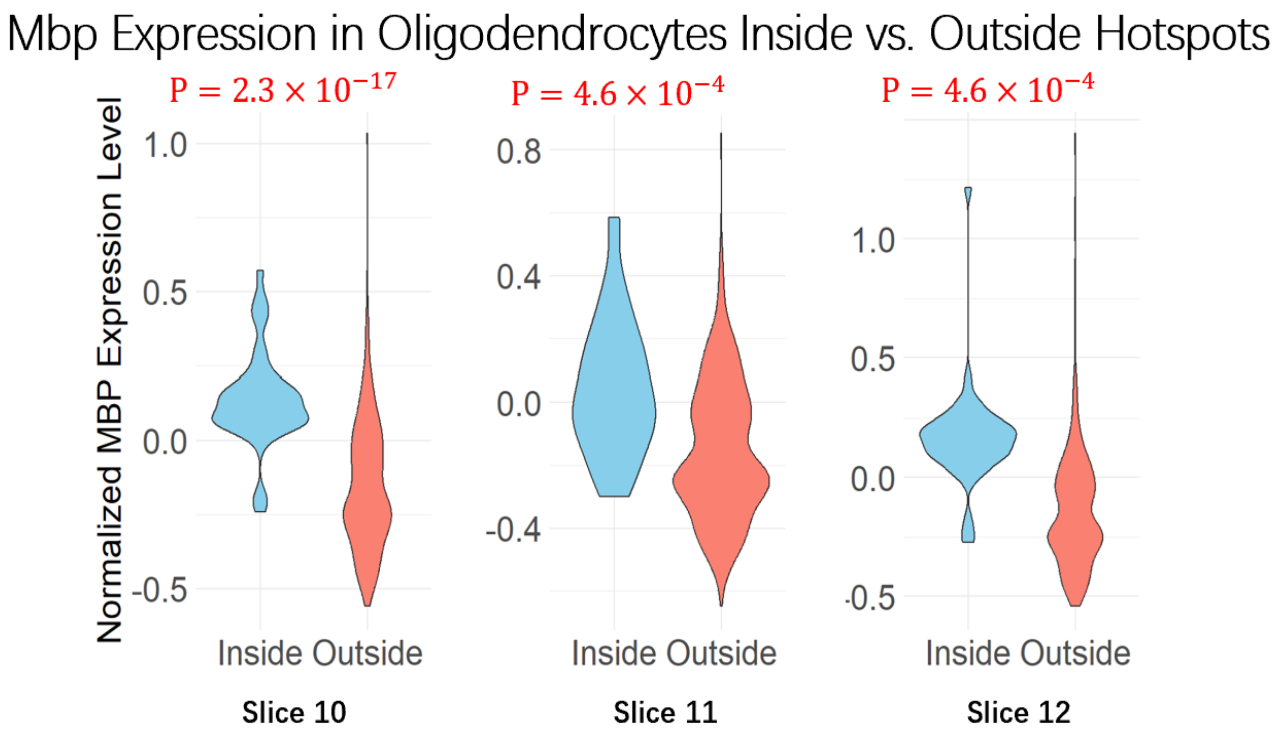

**Figure S19: The expression level of Mbp is consistently higher in oligodendrocytes within the Oligo-Oligo hotspot, compared to those outside the Oligo-Oligo hotspots in slices 10-12.**

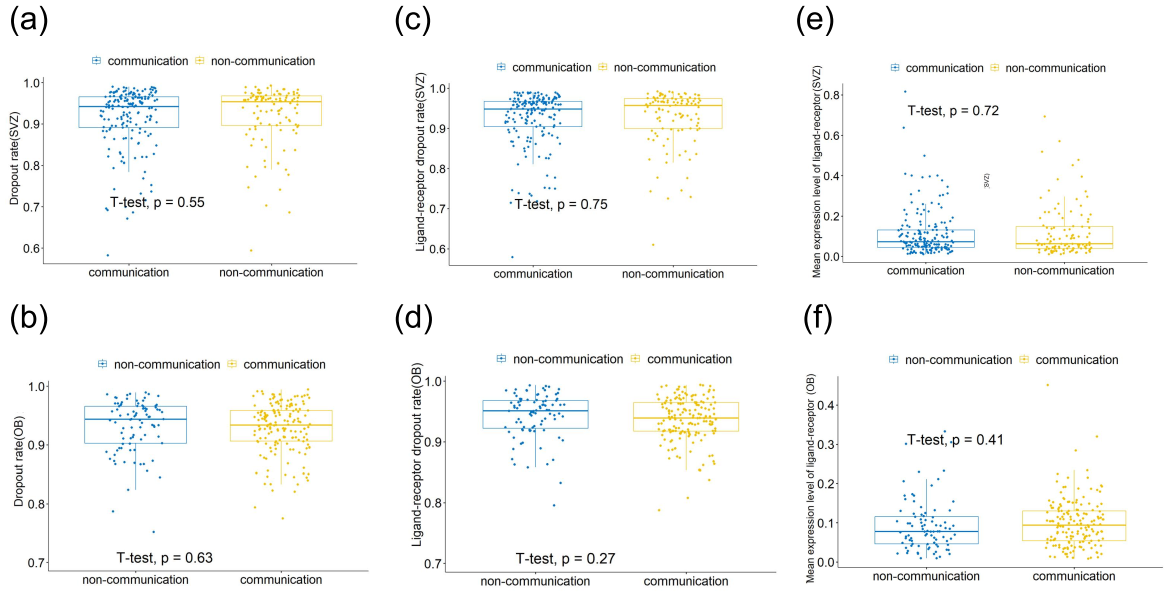

**Figure S20: The dropout rate and mean expression levels of ligands and receptors are comparable in cells involved and not involved in communication. Boxplot of gene dropout rate and mean expression level in the communication group and the non-communication group. (a),(b): Gene dropout rate for each cells in different slice. (a): SVZ (b): Olfactory bulb. (c),(d):Ligand-receptor dropout rate for each cells in different slice. (c):SVZ (d):Olfactory bulb. (e),(f): The mean expression level of ligand-receptor for each cells. (e):SVZ (f):Olfactory bulb.**
